# An active matter model captures the spatial dynamics of actomyosin oscillations during morphogenesis

**DOI:** 10.1101/2024.10.04.616649

**Authors:** Euan D. Mackay, Aimee Bebbington, Jens Januschke, Jochen Kursawe, Marcus Bischoff, Rastko Sknepnek

## Abstract

The apicomedial actomyosin network is crucial for generating mechanical forces in cells. Oscillatory behaviour of this contractile network is commonly observed before or during significant morphogenetic events. For instance, during the development of the *Drosophila* adult abdominal epidermis, larval epithelial cells (LECs) undergo pulsed contractions before being replaced by histoblasts. These contractions involve the formation of contracted regions of concentrated actin and myosin. However, the emergence and control of pulsed contractions are not fully understood. Here, we combined in vivo 4D microscopy with numerical simulations of an active elastomer model applied to realistic cell geometries and boundary conditions to study LEC actomyosin dynamics. The active elastomer model was able to reproduce in vivo observations quantitatively. We also characterised the relationship between cell shape, cell polarity, and actomyosin network parameters with the spatiotemporal characteristics of the contractile network both in vivo and in simulations. Our results show that cell geometry, accompanied by boundary conditions which reflect the cells’ polarity, is essential to understand the dynamics of the apicomedial actomyosin network. Moreover, our findings support the notion that spatiotemporal oscillatory behaviour of the actomyosin network is an emergent property of the actomyosin network, rather than driven by upstream signalling.

## I. INTRODUCTION

Proper execution of morphogenesis requires cells to coordinate their behaviours at the tissue scale. This coordination occurs via cellular processes that are shared across many organisms [1]. For example, cell rearrangements, cell migration and cell shape changes drive developmental processes, such as gastrulation, neurulation and axis elongation, which have been extensively studied in model organisms such as *Drosophila* [2], chicken [3], *Xenopus* [4] and zebrafish [5]. Cell rearrangements and cell shape changes, such as apical constriction, require cells to generate, transmit, and exert mechanical forces on each other and their surroundings [6]. Understanding the mechanistic basis of force generation will be crucial to gaining insights into diseases caused by defective cell behaviour, such as neural tube disorders [7]. Moreover, cell mechanics are known to underlie hallmarks of cancer [8]. Most of the force generation during morphogenesis originates in the actomyosin cytoskeleton [2, 3, 9–12]. The contraction of the actomyosin cytoskeleton is due to the activation of the motor protein non-muscle myosin II, which forms minifilaments that crosslink actin filaments and move them relative to each other [13]. Therefore, to understand collective cell behaviours, we must investigate how the cytoskeleton generates and transmits mechanical forces.

The cytoskeleton is an inherently dynamic structure, which is particularly evident from the frequent observation of periodic, spatiotemporal contractions and expansions. These have been described as pulsations, flows, pulsed contractions, or oscillations [15, 16]. Such pulsations have been seen in a variety of invertebrate model systems, e.g. in *C. elegans* embryos [17], in starfish oocytes [18], and in various processes in the *Drosophila* embryo, e.g. during gastrulation [19], dorsal closure [20], germband extension [21], eye morphogenesis [22], and oocyte formation [23]. Moreover, pulsations have been observed in vertebrates, e.g. during *Xenopus* neurulation [4], mouse fertilisation [24], and mouse compaction [25], as well as in a variety of in vitro cell cultures [26].

Polar active gel theories that model the actomyosin network as a viscoelastic fluid of interconnected polar actin filaments locally contracted by myosin molecular motors have been successful in describing many aspects of the cytoskeleton mechanics [27, 28]. A version of this approach that treats the actomyosin network as an isotropic active elastomer with a short time scale viscous behaviour has also proven effective in descring actomyosin oscillations [29, 30]. However, despite recent insights into the emergence of actomyosin oscillations [11, 31, 32], a largely unanswered question is how subcellular actomyosin contractions are spatially organised and how cell size and shape influence the contractile dynamics of the cytoskeletal network.

Discussions are also still ongoing regarding the origin of the pulsations, contrasting emergence versus upstream signalling [15, 33]. In favour of pulsations being emergent, it has been argued that advection, caused by myosin-induced contractions, can by itself produce enough of a positive feedback mechanism to sustain actomyosin oscillations [11, 31]. On the other hand, several studies suggest that the pulsations are largely driven by upstream signalling mechanisms, such as rhythmical activation by the small Rho GTPase Rho1/RhoA, which activates Rho kinase (Rok), which in turn activates Myosin-II [34–37]. Rhythmical activity of Rho1 is generated by its consecutive activation and deactivation by Rho guanine nucleotide exchange factors (RhoGEFs) and Rho GTPase activating proteins (RhoGAPs), respectively [38–40].

Here, we combine in vivo 4D microscopy and an active elastomer model to investigate the mechanism of actomyosin pulsations of *Drosophila* larval epithelial cells (LECs). To make direct comparisons between the model and experiments, we imaged LECs in high spatial and temporal resolution and used the finite element method to solve the active elastomer model on realistic cellular geometries with several different boundary conditions.

Due to their large size (up to 70 μm), LECs are well-suited to study subcellular actomyosin dynamics. During morphogenesis of the adult abdominal epidermis, the large polyploid LECs are replaced by diploid histoblasts, which form the adult epidermis (Fig. 1a) [41, 42]. During this process, LECs undergo directed migrations and apical constriction before they delaminate and die [43]. Migrating LECs create a crescent-shaped lamellipodium in the direction of movement (Fig. 1b) [43]. During their migration and constriction, cells exhibit pulsed contractions, enabling the study of actomyosin oscillations in different contexts [14]. Migrating LECs display a planar cell polarity with the lamellipodium at the front and contractile activity in the back, whereas constricting LECs exhibit a radial cell polarity with contractile activity in the cell centre (Fig. 1b) [14]. The behaviour of the actomyosin network in LECs, as well as in gastrulating *Drosophila* cells, depends on cellular contractility levels, i.e. myosin activity, with pulsed contractions only taking place with intermediate levels of actomyosin activity [11, 14].

**Figure 1.**
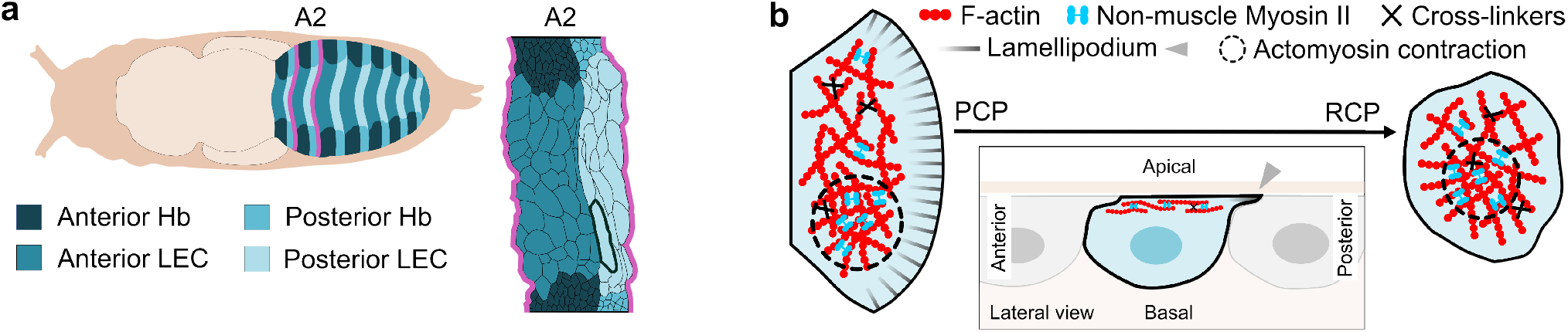
Larval epithelial cells (LECs) are replaced by histoblasts during *Drosophila* abdominal morphogenesis. **a**. Schematic of a pupa (left) and zoom-in of abdominal segment A2 (right). Dorsal view, anterior is to the left. Histoblasts (Hb) replace the larger LECs. Cells of the anterior (dark) and posterior (light) compartments are indicated by shade. We focus our analysis on LECs at the anterior edge of the posterior compartment of A2, that are not in direct contact with Hbs (bold outline). **b**. Schematic of a migrating and constricting LEC. The apicomedial network is a thin, crosslinked network of filamentous actin underlying the apical cell membrane, on which non-muscle Myosin-II acts to generate contractile stress. Nuclei are positioned below the apicomedial network (ellipses in lateral view). Migrating LEC shows planar cell polarity (PCP), with a lamellipodium at the front and the contractile network in the back; constricting LEC shows radial cell polarity (RCP), with the contractile network in the cell centre [14].

Our approach reveals that realistic cell shapes accompanied by the appropriate boundary conditions, which reflect the planar and radial cell polarities of the LECs, respectively, are crucial to understanding the nature of the observed actomyosin pulsations. Moreover, our results support the notion that pulsed contractions are an emergent property of the actomyosin network, rather than driven by upstream signalling networks.

## II. RESULTS

### A. LECs exhibit distinct patterns of apicomedial actomyosin activity

To characterise the spatial, sub-cellular dynamics of actomyosin contractions in LECs, we imaged cells in high spatial and temporal resolution. We focused our analysis on dorsal LECs of segment A2 at the anterior edge of the posterior compartment that are not in direct contact with histoblasts to ensure comparability between cells and with previous work (Fig. 1a) [14]. We labelled the apicomedial actin network with the F-actin marker GMA-GFP [44]. The distribution of F-actin within the apicomedial network is a good proxy for that of myosin II (Supplementary Fig. S1) [14]. Visualising the apicomedial actin network approximately every three seconds allowed us to characterise actomyosin contractions in detail (Fig. 2).

**Figure 2.**
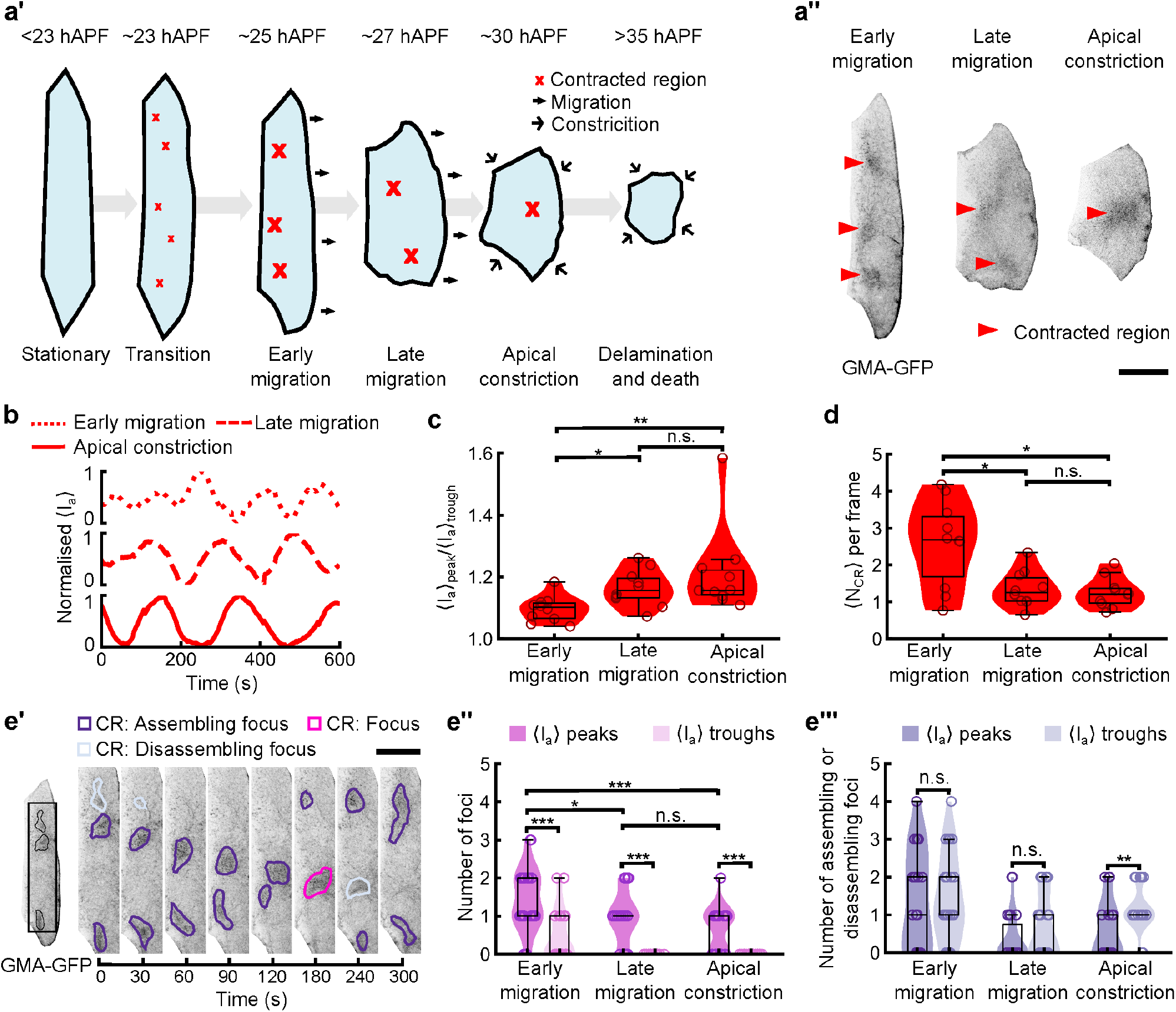
Actomyosin oscillations in LECs during abdominal morphogenesis. **a’**. Schematic of the typical behaviour of LECs during abdominal morphogenesis. Small, transient contractions (small red crosses) appear in the apicomedial actomyosin networks of stationary LECs when histoblasts begin to proliferate. Then, LECs establish lamellipodia and actively migrate (black arrows). Eventually, LECs transition into constrictive behaviour (black chevrons), before delaminating and dying. Distinct “contracted regions” (CRs) (large red crosses) are present in the apicomedial actomyosin networks of migrating and constricting LECs. **a’’**. LECs of the three pulsatile stages. F-actin labelled with GMA-GFP. CRs are visible (red arrowheads). Scale bar: 20 μm. **b**. LECs show actomyosin oscillations. Representative time series of normalised spatially averaged apical fluorescence intensity, ⟨*I*_a_⟩, of early and late migrating, and apically constricting LECs expressing GMA-GFP. **c**. The ratio of peak to trough spatially averaged apical GMA-GFP fluorescence intensity increases significantly between early and late migration. **d**. The mean number of CRs, ⟨*N*_CR_⟩, detected in each frame decreases between early and late migration. **e**. Studying focus assembly and disassembly. **e’**. CRs (purple) propagate through the apicomedial network before coalescing into foci (magenta) and subsequently disassembling (lavender). Scale bar: 10 μm. **e’’, e’’’**. Comparison of numbers of distinct foci (**e’’**) and assembling or disassembling foci (**e’’’**) present at peaks and troughs in spatially averaged apical GMA-GFP fluorescence intensity time series across the three pulsatile LEC stages. Early migration *n* = 29 peaks, 29 troughs; late migration *n* = 26 peaks, 26 troughs; apical constriction *n* = 27 peaks, 28 troughs. \**p <* 0.05, ** *p <* 0.01, \*\*\**p <* 0.001. In each panel, *n* = 10 recordings per stage. Pairwise significance comparisons were done with a two-sided Mann-Whitney test.

At the beginning of abdominal morphogenesis, the LECs are stationary until approximately 23 hours after puparium formation (APF), during which time no contractions are visible. This is followed by a fluctuation phase in which numerous small, transient contractions appear, accompanied by small fluctuations in apical cell shape. By about 25 h APF, LECs establish lamellipodia and begin to migrate posteriorly [43]. In this early migration phase, LECs display distinct regions of higher actin density, which we refer to as “contracted regions” (CRs) (Fig. 2a’). A useful measure is the time-averaged number of CRs per cell, ⟨*N*_CR_⟩. CRs typically appear in peripheral locations before propagating through the network, coalescing into high fluorescence intensity “foci”, and subsequently disassembling. These oscillatory patterns persist through late migration (∼27 h APF) and into apical constriction (∼ 30 h APF) (Fig. 2a‘‘, Supplementary Video 1). Overall, this dynamic activity results in oscillations in spatially averaged GMA-GFP fluorescence intensity in early migration, late migration, and apical constriction (Fig. 2b). The amplitude of these oscillations increases slightly between early and late migration (Fig. 2c).

While the qualitative dynamics of individual CRs, i.e. initiation, propagation, focus formation, and disassembly, are similar, early migrating LECs display more CRs (2.6 ± 1.1) than either late migrating or apically constricting cells (1.3 ± 0.4 and 1.4 ± 0.5, respectively) (Fig. 2d).

To investigate whether the different number of CRs in the different pulsatile LEC stages reflects a difference in the spatiotemporal behaviour of the actomyosin network, we investigated how CRs coalesce into foci (Supplementary Video 2). To this end, we located peaks and troughs in the spatially averaged fluorescence intensity time series and classified the CRs detected at each timepoint as being either a stable focus or belonging to an assembling or disassembling focus (Fig. 2e‘). We found that LECs in late migration and apical constriction showed 0.9±0.6 and 0.6±0.5 foci at intensity peaks, respectively. There were no foci at intensity troughs (Fig. 2e‘‘), with 58% of troughs in late migration, and 7% in apical constriction, showing a complete absence of contracted regions (Fig. 2e‘‘‘). LECs in early migration, on the other hand, showed 1.4±0.8 foci at intensity peaks and 0.5±0.7 foci at intensity troughs (Fig. 2e‘‘), with similar numbers of assembling/disassembling foci at peaks and troughs (Fig. 2e‘‘‘). Thus, during late migration and constriction, CRs tended to coalesce into one focus per oscillation peak. This observation fits the assumption that oscillations in averaged fluorescence intensity are caused by the rhythmical contraction of the actomyosin network coalescing into a single focus. Early migrating LECs, on the other hand, support multiple foci in different stages of assembly (Fig. 2e‘‘), with oscillations in averaged fluorescence intensity arising from the combined activity of those foci.

The subcellular spatiotemporal patterns of pulsed contractions not only depend on actomyosin activity but also correlate with the cells’ aspect ratio [14]. To gain further insights into the nature of LEC shape change, we quantified cell shapes by measuring the maximum and minimum Feret diameters, i.e. the maximum and minimum distances between two parallel lines tangent to the contour of the cell boundary (Supplementary Fig. S2a). Early migratory LECs are dorsoventrally elongated, with an aspect ratio of ≈ 5 (*n* = 10 recordings). From early to late migration, mean apical area remains similar (1400±500 μm^2^ to 1500±500 μm^2^), but aspect ratio decreases significantly to ≲ 3, i.e. cells begin to round up (Supplementary Fig. S2a‘‘,b). The transition from late migration to constriction leads to a further rounding up of the cells, involving a reduction in average apical area (to 900±300 μm^2^) and a decrease of the aspect ratio to ≲ 2 (Supplementary Fig. S2 a’’,b) (*n* = 10 recordings). It is unclear how the described diverse, yet rather regular patterns of actomyosin oscillations emerge mechanistically. To untangle the potential effects of apical cell shape, cell polarity, and contractility, we utilised a twodimensional active elastomer model of the apicomedial actomyosin network.

### B. Oscillations in the apicomedial actomyosin network can be described by a two-dimensional active elastomer model

At the time scale of a pulsed contraction the actomyosin network in LECs can be modelled as a twodimensional active elastomer [29–32], with the overdamped dynamics described by

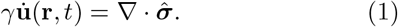

The overdot is the time derivative and *γ* is the friction constant, related to the viscosity of the permeating fluid and the typical distance between actin filaments [45]. **u**(**r**, *t*) is the displacement from a stress-free rest configuration at the position **r** and time *t*, and 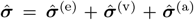 is the total mechanical stress, where 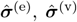, and 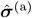 are elastic, viscous, and active stresses, respectively, meaning the constiuitive relationship is that of the Kelvin-Voight model. Expression for 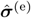 and 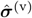 are derived assuming small strains and strain rates as well as an isotropic and homogeneous network (Materials and Methods).

Active stresses are generated by myosin [46, 47] and can be modelled as [31]

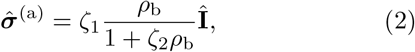

where *ρ*_b_ a scalar density field of non-muscle myosin II motors bound to actin filaments, and *ζ*_1_ and *ζ*_2_ are constants. For the network to be contractile *ζ*_1_, *ζ*_2_ *>* 0 [31]. A cartoon of the key ingredients of the model is sketched in Supplementary Fig. S3.

The bound myosin is both advected with the network and is diffusing relative to it with diffusion constant *D*. These two processes are described by an advectiondiffusion equation,

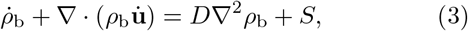

where *S* is a turnover of the bound myosin specified by binding and unbinding rates *k*_b_ and *k*_u_, respectively. Single-molecule experiments reported *k*_b_ ∼ 10 s^−1^ [48] and found strain-dependent unbinding rates [48, 49]. Therefore, we set 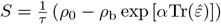 where 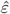 is the strain tensor, with 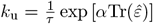 and 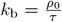, where *ρ*_0_ is the equilibrium density of bound myosin. *τ* controls the intrinsic timescale associated with myosin turnover. *α >* 0 (*α <* 0) represents positive (negative) feedback where an increased compression results in slower (faster) myosin unbinding. We set *α* = −1 and define strain-free myosin turnover rate 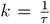. We can, thus, track only *ρ*_b_ and ignore the unbound myosin [31]. Finally, we chose 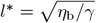 as the unit of length, *t*^*^ = (*l*^*^)^2^*/D* as the unit of time, and *ρ*^*^ = 1*/ζ*_2_ as the unit of density. The parameter *ζ* controls the magnitude of active stress and is the dimensionless analogue of parameter *ζ*_1_ in Eq. (2). The non-dimensional equations are shown in Materials and Methods, and model parameters and their values are summarised in Supplementary Tab. S1.

For the initial conditions, we used **u**(**r**, *t* = 0) = *δ***u**(**r**) and *ρ*_b_(**r**, *t* = 0) = *ρ*_0_ + *δρ*(**r**), where *ρ*_0_ is a constant, and *δ***u**(**r**) and *δρ*(**r**) are small local random perturbations with zero mean and standard deviation of 0.005.

Depending on stage of the LECs life cycle, we considered three distinct types of boundary conditions for *ρ*_b_: 1) *no flux* condition applied everywhere (used for radially polarised cells in the constriction phase), and defined as 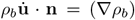 on *∂*Ω, where **n** is the unit-length normal vector to the cell boundary *∂*Ω. 2) *Strong mixed* boundary conditions (used for planar polarised cells in the late migration phase), where the side of the cell without lamellipodia have no-flux boundary conditions and the side with lamellipodium has myosin fixed at *ρ*_b_ = *ρ*_0_ by a Dirichlet boundary condition. 3) *Weak mixed* boundary conditions that are same as 2) but with myosin on the lamellipodium fixed at 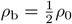 (used for planar polarised cells in the early migration phase). Although lamellipodia are dynamic structures, they can be assumed static at time scales of interest. Their presence prevents contractility in the front of the cell [14], which justifies fixing *ρ*_b_ at the cell boundary with a lamellipodium.

We also assumed that, at time scales of interest, cell boundaries are pinned, meaning there is no displacement of the cell boundary itself, which corresponds to a Dirichlet boundary condition on **u**, i.e. **u** = **0** on *∂*Ω. This assumption significantly simplifies the numerical simulations. The combination of a no flux boundary condition on myosin and a pinned condition on **u** is known as a *natural* boundary condition. For comparison with existing literature, e.g. Refs. [29–32], we also considered periodic boundary conditions, but only in simple rectangular geometries.

### C. Active elastomer dynamics in rectangular geometries partly capture observed behaviours

We simulated dynamics by solving the model equations with the finite element method (FEM) implemented in FEniCS [50, 51] (Materials and Methods). The advantage of using FEM is that it is possible to study realistic domain shapes. However, for simplicity, we first studied simple rectangular geometries (of size 20*l*^*^ × 8*l*^*^) using both no-flux and periodic boundary conditions. In particular, we explored the effects of tuning the strain-free myosin turnover rate, *k*, and the actomyosin activity, *ζ*. For each of these we analysed the state of the system at time *t* = 500 *t*^*^ after the steady state was reached (see Supplementary Fig. S4a).

For no-flux boundary conditions, we observed two distinct steady states (Fig. 3a). At low actomyosin activities or high myosin turnover rates, the system relaxed back into the homogeneous state. At higher activities or lower turnover rates, the system developed a complex spatiotemporal pattern of myosin density (Fig. 3c; Supplementary Videos 3,4). These regions are highly reminiscent of CRs seen in the LECs. We, however, note that the behaviour for *k* = 0 line in the phase diagram is noticeably different, because in this instance the system becomes mass conserving, leading to different dynamics (Supplementary Information Sec. S6, Supplementary Fig. S5).

**Figure 3.**
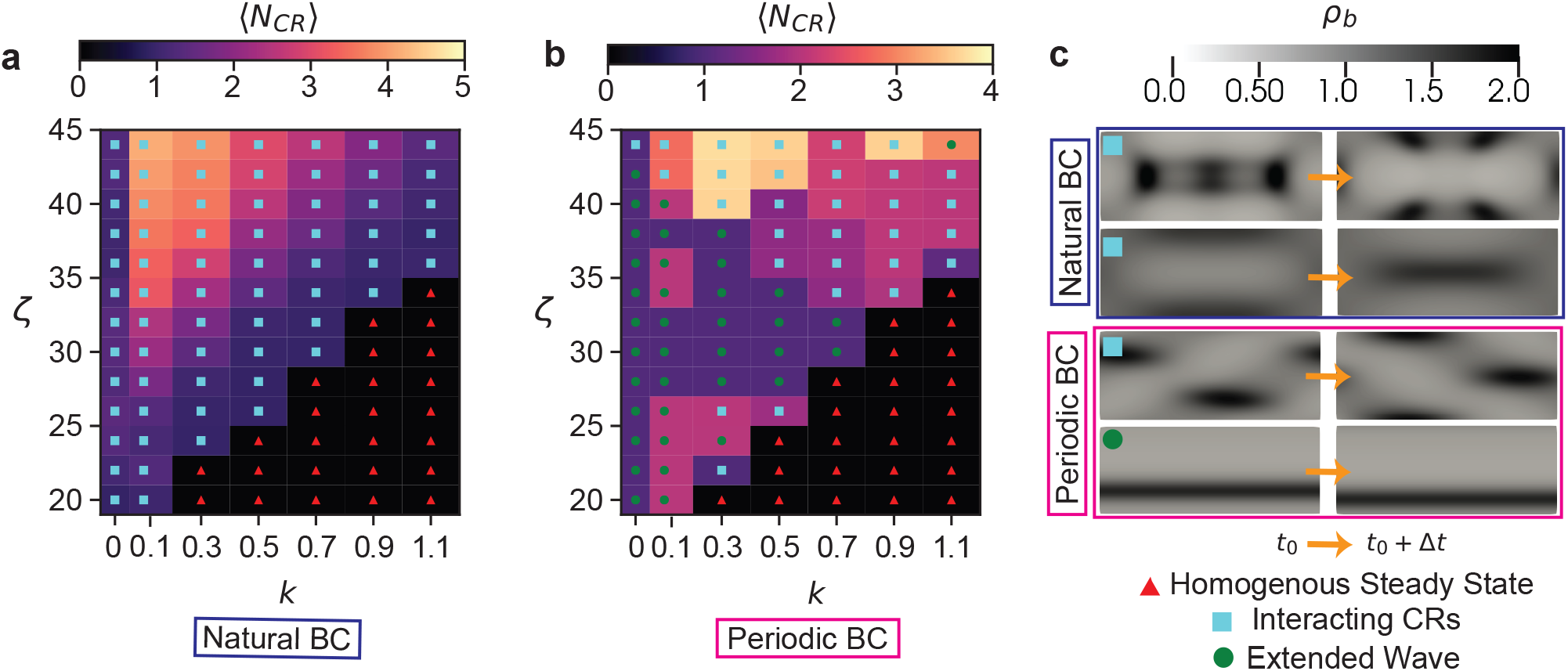
The overall behaviour of the active elastomer as a function of model parameters. **a**. Phase diagram with no-flux (natural) myosin boundary conditions (BC), as a function of activity *ζ* and strain-free myosin turnover rate 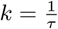, where *τ* is the myosin turnover time. We observed two distinct states: 1) a homogeneous steady state with **u** = **0** and *ρ*_*b*_ = *ρ*_0_ (red triangles), and 2) one or more interacting CRs exhibiting complex spatiotemporal patterns (cyan squares). **b**. Phase diagram of the system with periodic boundary conditions. Here, we observed a third state, where the CRs merge and form a moving, phase-separated state (extended wave, green circle). **c**. Examples of the behaviour of the system showing snapshots of bound myosin density (taken Δ*t* ∈ {0.75*t*^*\**^, 0.6*t*^*\**^, 0.4*t*^*\**^, 0.25*t*^*\**^} apart for the four snapshot pairs). Only interacting CRs and extended wave states are shown. With no-flux boundary conditions, close to the homogeneous steady state, we typically see a single extended CR (top snapshot). As *ζ* is increased (or *k* is decreased), more CRs appear (second snapshot). For periodic boundaries, we saw a similar state with multiple CRs, although their arrangement appeared to be a bit more regular (third snapshot). We also saw a new state where the CRs all merge into one large CR, extended in one direction (bottom snapshot). All phase diagrams simulations were performed on a rectangular mesh of size 20*l*^*\**^×8*l*^*\**^ with all non-specified parameter values given in Supplementary Tab. S1. In both phase diagrams, colour bars represent the time average of the number of contracted regions, ⟨*N*_CR_⟩. For discussion about subtleties with setting *k* = 0 (i.e. *τ* → ∞) see Supplementary Information Sec. S6 and Supplementary Fig S5.

To compare to previous work [31], we also constructed a phase diagram of the system with periodic boundary conditions (Fig. 3b), where we observed the same two states as in the no-flux case (Fig. 3a). The CRs were, however, more regular and periodic (Fig. 3c; Supplementary Video 5), which is likely due to self-interactions across periodic boundaries. In addition, a third state emerged, where the CRs coalesce into a single, myosin-rich linear domain of finite width (Fig. 3c; Supplementary Video 6). Since we only observed this state with periodic boundaries, it is plausible that it is an artefact caused by self-interaction across the boundaries. These findings are broadly consistent with the one-dimensional analysis of a similar model discussed in Ref. [31], with an important difference that we did not observe the inhomogeneous steady state since in our model active stresses are strain-independent.

For the system with no-flux boundary conditions, we next explored the behaviour as a function of activity *ζ* and system size (Supplementary Fig. S4b). We kept the length of the short side of the rectangle constant and varied the length of the long side, *L*_long_, to mimic the shape changes LECs undergo (Supplementary Fig. S2a’’). Once again, at low *ζ*, the system was in the homogeneous steady state, and as *ζ* increased, the system developed CRs, which increased in number with increasing *ζ*. We observed a similar effect as *L*_long_ was increased, with no CRs observed at very low *L*_long_, and their number increasing with *L*_long_. This suggests that CRs have a characteristic minimum size *ℓ*_CR_, which depends on activity *ζ*. Consequently, for a given value of *ζ* when *L*_long_ *< ℓ*_CR_, we expect to observe no CRs.

Finally, we also varied the equilibrium density of bound myosin, *ρ*_0_, and activity *ζ*. We found that at low values of *ρ*_0_, the transition between the two observed states happens gradually, with the number of simultaneous CRs gradually increasing with *ζ*. However, this transition became sharper for higher values of *ρ*_0_, with the rapid onset of a state with up to five CRs, at large values of *ζ* (Supplementary Fig. S4c).

In summary, we found that for a suitable set of parameters, the model creates oscillating CRs with both periodic and no-flux boundary conditions. The precise number and nature of these CRs depends on parameter values, as well as the boundary conditions. We also captured the homogeneous to oscillatory steady state transition using linear stability analysis (Supplementary Information Sec. S7 and Supplementary Fig. S6). However, linear stability analysis is unable to capture all features observed in experiments and simulations, as recently discussed for a related model [52].

### D. The active elastomer model captures spatial dynamics of apicomedial pulsation in realistic cell geometries

To compare the model with in vivo data, we performed simulations using meshes constructed from real cell geometries extracted from images of LECs in early migration, late migration and apical constriction stages. We compared the dynamics of the measured in vivo apical F-actin fluorescence intensity and simulated myosin density fields, as well as measured and modelled velocity fields. We recall that apical F-actin fluorescence intensity (GMA-GFP) is an experimental proxy for myosin density (Supplementary Fig. S1). We extracted the experimental flow field velocity from the imaging data using optical flow analysis [53] (Materials and Methods), while in the model, we computed it as 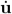. We found that the model qualitatively agreed with observed dynamics, e.g. CRs initiated in approximately the same locations, flows were typically in the same directions with similar numbers of CRs present at any one time (Fig. 4 – left column). Furthermore, spatially averaged oscillations of the F-actin or myosin for both the model and experiments were approximately out of phase with those of the divergence of the velocity field, ∇ · **v**, a quantity that captures local contractions and expansions (Fig. 4 – right column).

**Figure 4.**
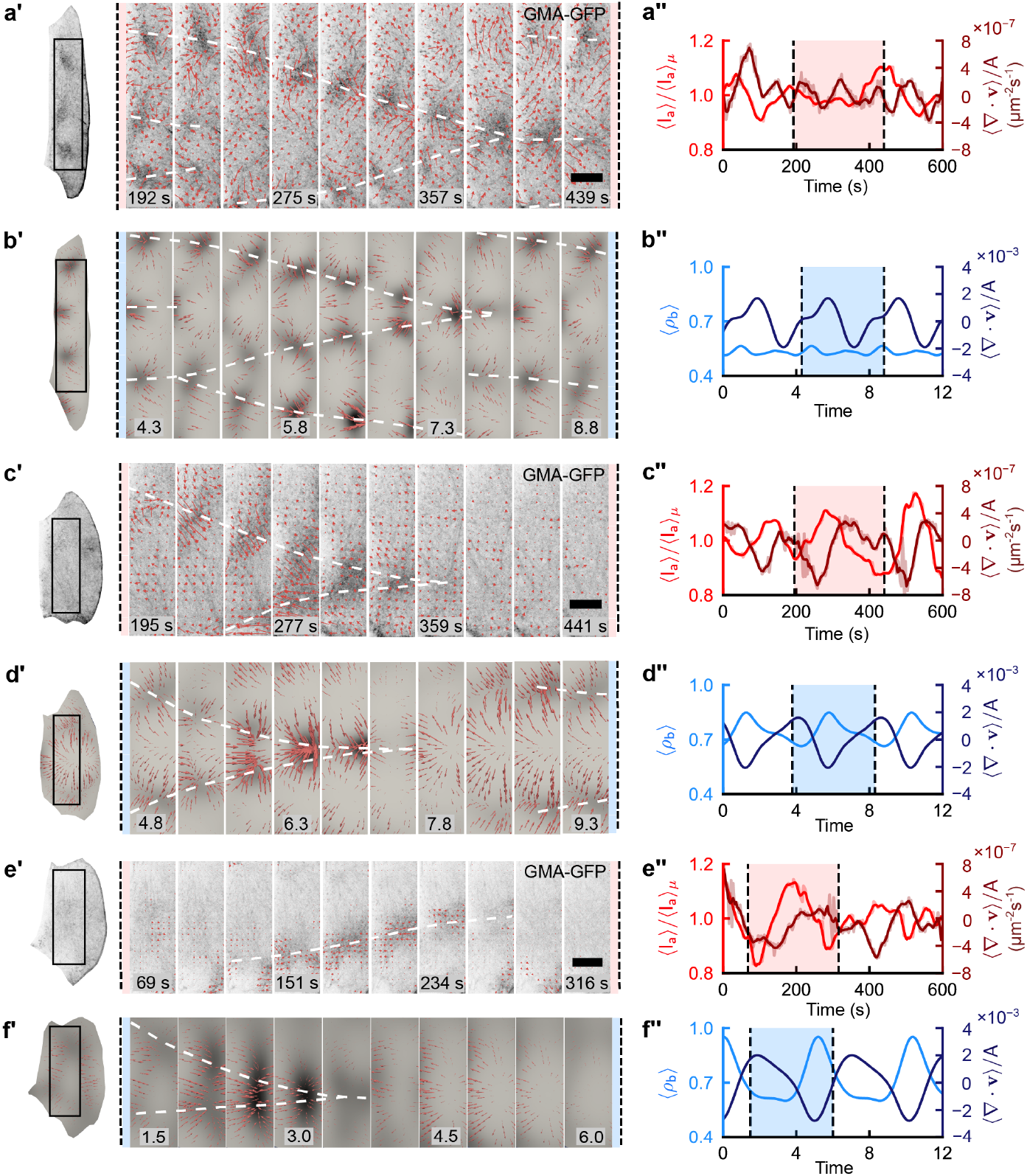
Simulations on in vivo cell shapes qualitatively capture the number, location, and movement of CRs observed in vivo. The left column shows cellular subregions (black rectangles) of in vivo F-actin fluorescence (**a’**,**c’**,**e’**) and modelled myosin density (**b’**,**d’**,**f’**) distributions in space, as a set of evenly spaced time points. Early migrating (**a’**,**b’**), late migrating (**c’**,**d’**) and constricting cells (**e’**,**f’**) shown. Shading corresponds to fluorescence intensity and myosin density, respectively. Red arrows denote the velocity field of the actomyosin network. Since it is not possible to directly compare numerical values of measured and computed velocity fields, for clarity we scaled velocity vectors obtained from the model by a factor of 0.5. Both in the in vivo experimental and model data, CRs are forming and flow through the cell (white dashed lines are to guide the eye). The right column shows time series of spatially averaged F-actin fluorescence intensity, normalised by the mean (**a’’**,**c’’**,**e’’**), and myosin density (**b’’**,**d’’**,**f’’**) over the whole cell (light colour curves). Early migrating (**a’’**,**b’’**), late migrating (**c’’**,**d’’**), and constricting cells (**e’’**,**f’’**) shown. We also plot the spatially averaged divergence per unit area of the velocity field (dark colour curves). Both in the in vivo and in the model data, oscillations are visible, which contract and expand. For imaging data, denoising was performed (see Materials and Methods). Denoised data is represented with solid lines while raw data is shown as faded lines. Shaded regions between vertical dashed lines correspond to the time sequence shown in the left column. Scale bars: 10 μm. Videos of the images and model data are shown in Supplementary Videos 7-12.

We obtained an excellent qualitative agreement between the experimental and numerical data by tuning only the activity *ζ* and changing the boundary conditions while keeping all other parameters fixed. We found the best agreement between experiments and the model for *ζ* = 40 (early migration), *ζ* = 36 (late migration), and *ζ* = 30 (apical constriction). This suggests that as replacement of LECs proceeds, their level of active contractile stress decreases.

Additionally, we found that cells in early migration matched the data best when weak mixed boundary conditions were applied. In contrast, cells in late migration required strong mixed boundary conditions and apically constricting cells required no flux conditions. This is consistent with the motivation for varying the boundary conditions in different stages of the LEC life cycle, proposed above. Migrating LECs are planar polarised, generating a lamellipodium at their front, while apically constricting LECs appear radially polarised [14]. It is, therefore, reasonable to assume that the effect of the cellular asymmetry first increases over time as cells commit to migration, and then disappears in the transition to apical constriction, when the lamellipodium disappears.

Overall, our model supports the notion that both actomyosin activity and cell polarity affect the behaviour of the actomyosin network. Moreover, our data show that it is helpful to use realistic cell geometries for the model to match the in vivo data. This indicates that cell shape also affects actomyosin behaviour.

### E. Actomyosin activity, cell shape, and cell polarity influence the spatiotemporal behaviour of the actomyosin network

To gain further insights into the effects of actomyosin activity, cell shape, and cell polarity on the spatiotemporal behaviour of the actomyosin network, we quantitatively compared each experimental recording with numerical simulations using in vivo cell shapes. We focused on features of the actomyosin network that change between early and late migration in vivo, namely the number of CRs per cell (Fig. 2d) and the number of CRs present at peaks in spatially averaged GMA-GFP intensity (Fig. 2e’’). Furthermore, we investigated the subcellular CR initiation sites and oscillation period. We initially used model parameters that resulted in the best agreement between experiments and simulations. We found that for all analysed features, the model captured the trends observed in vivo (Fig. 5).

**Figure 5.**
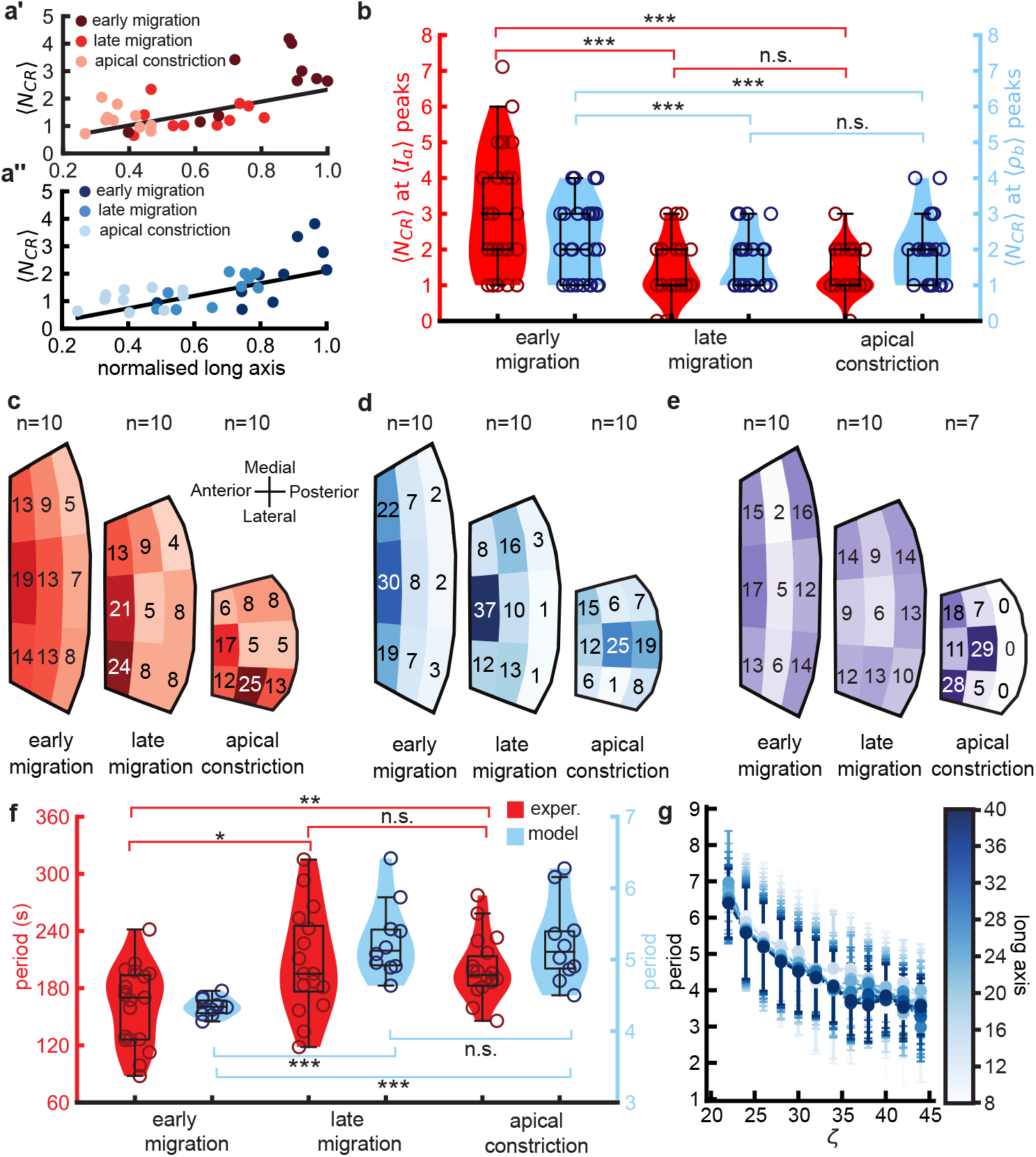
Quantitative comparison of experimental data with simulations for in vivo cell shapes. **a’**,**a’’**. Mean number of CRs vs. long axis length (normalized to the greatest long axis length) in experiments (**a’**) and the model (**a’’**); *n* = 10, all cases. Error bars not shown for clarity; the standard deviation is typically ∼ 1.5. **b**. Number of CRs at spatially averaged intensity maxima for the three LEC stages (red – experiments, *n* = 29; early migration, *n* = 26; late migration, *n* = 27; apical constriction; blue – model, *n* = 275; early migration, *n* = 249; late migration, *n* = 229 in apical constriction). Every tenth model data point is shown. **c**. Distribution of CR initiation sites in the three LEC stages in experiments. *n* = 10 in all cases. Colour shade is proportional to the percentage of initiation sites in each cell segment. **d**. Same as (**c**) for the model with weak mixed (early migration), strong mixed (late migration), and no flux (apical constriction) boundary conditions. *n* = 10 in all cases. **e**. Same as (**d**) with swapped boundary conditions, i.e. no flux (early and late migration), strong mixed (apical constriction). *n* = 10 in early and late migration, *n* = 7 in apical constriction. For each experiment and simulation, initiation sites were located within the appropriate cell segment (Materials and Methods), and percentages computed and averaged over all *n* instances. **f**. Period of spatially averaged actin fluorescence intensity (experiments – red, *n* = 19, early migration; *n* = 16, late migration; *n* = 18, apical constriction) and myosin density (model – blue, *n* = 10, all stages) in the three LEC stages. **g**. Period of oscillations vs. activity *ζ* in the model on rectangular domains for varying lengths of the long axis and fixed short axis (cf. Supplementary Fig. S4b). Error bars show standard deviation (across space) of dominant period. \**p <* 0.05, ** *p <* 0.01, \*\*\**p <* 0.001. Pairwise significance comparisons were done with a two-sided Mann-Whitney test.

#### 1. The number of CRs depends on actomyosin activity, cell shape and cell polarity

We observed a significant positive correlation between the time averaged number of simultaneously detected CRs (⟨*N*_CR_⟩) and cells’ long axis length in both experiments (Spearman *r* = 0.58, *p* = 0.001) and model simulations (Spearman *r* = 0.68, *p* = 0.0002) (Fig. 5a’,a’’). This is consistent with the observation that, for simulations on rectangular domains, ⟨*N*_CR_⟩ increases with both domain size and actomyosin activity *ζ* (Fig. 3a,b and Supplementary Fig. S4b,c).

To further investigate the effects of cell contractility, cell polarity, and cell shape on the number of CRs, we calculated ⟨*N*_CR_⟩ using several different model scenarios (Supplementary Information Sec. S8; Supplementary Fig. S7). We found that keeping the actomyosin activity constant (*ζ*=36) and applying variable boundary conditions for the three stages (weak mixed for early migration, strong mixed for late migration, no flux for constricting cells) resulted in an insignificant correlation between ⟨*N*_CR_⟩ and cell length (Spearman *r* = 0.36, *p* = 0.06) (Supplementary Fig. S7b). Applying no flux boundary conditions for each stage but adjusting actomyosin activities (*ζ* = 40 for early migration, *ζ* = 36 for late migration, *ζ* = 30 for constricting cells), resulted in a much stronger positive correlation (Spearman *r* = 0.86, *p* = 0.0002) (Supplementary Fig. S7c). Finally, if no flux boundary conditions and the same actomyosin activity (*ζ* = 36) were applied to all LEC stages, we found a slightly weaker, but still significant correlation (Supplementary Fig. S7d). Overall, the observed moderate positive correlation between ⟨*N*_CR_⟩ and cell length which was measured in vivo as well as in the model (Fig. 5a‘,a’’) appears to arise from the combined effects of contractility, cell size, and cell polarity, with increasing cell size and contractility acting to increase ⟨*N*_CR_⟩, while mixed boundary conditions make ⟨*N*_CR_⟩ more noisy compared to the no flux boundaries case.

Moreover, we found agreement between the number of CRs present at peaks in spatially averaged measured GMA-GFP intensity and model myosin density, with typically one or two CRs at peaks in late migration and apical constriction, and a greater number in early migration (Fig. 5b). Thus, during early migration, oscillations in averaged fluorescence intensity are not caused by the coordination of multiple CRs coalescing into a single focus, which would involve the coordinated contraction of the entire actomyosin network. Instead intensity maxima in early migration are a result of more complicated spatiotemporal organisation of many CRs. This behaviour is consistent with the reduction in oscillation amplitude observed in vivo (Fig. 2c) and in the model (Supplementary Information Sec. S9 and Supplementary Fig. S9), as when multiple oscillatory events are averaged over, interference effects lead to a reduction in amplitude.

#### 2. Cell polarity affects the location of CR initiation sites

To explore the spatial organisation of CRs, we mapped the locations where subcellular contractions start. In vivo, these initiation sites were predominantly distributed over the anterior side of migrating cells, whereas in constricting cells the anterior bias was much weaker. There was also a slight lateral bias (Fig. 5c) in the constricting cells. The model produced broadly similar results but with a sharper transition, and with no obvious anterior or lateral bias in constricting cells (Fig. 5d).

To investigate further, in simulations, we altered the boundary conditions. For early and late migrating cells, we switched to no flux, and for apically constricting cells, we used strong mixed boundary conditions. This resulted in the loss of anterior bias in the position of initiation sites in the two migrating stages, as well as the gain of anterior bias in the apically constricting cells (Fig. 5e). This indicates that the location of initiation sites is determined by cell polarity.

Polarisation could also explain the slight differences between the in vivo and modelling data, with the in vivo data having a small anterior and lateral bias during constriction (Fig. 5c), whereas this bias is absent in the modelling data (Fig. 5d). When LECs cease to migrate, they are typically approached by the histoblasts. This briefly polarises the LECs along the dorsoventral axis before they become radially polarised [14], which might bias the CR initiation sites laterally.

#### 3. The period of pulsed contractions is influenced by network contractility and cell polarity, but not cell shape

Next, we examined the oscillations in spatially averaged GMA-GFP fluorescence intensity and simulated myosin density that arise from the dynamics of CRs. In experiments, we found that the period of these oscillations increased by ≈30% from 161 ± 9 s (*n* = 19 cycles) during early migration to 210 ± 20 s (*n* = 16 cycles) in the late migration stage and remained approximately the same (201 ± 8 s, *n* = 18 cycles) in the apical constriction phase (Fig. 5f). The model is in good agreement with the experiments as simulations found spatially averaged periods of 4.3 ± 0.1 t^*^ (early migration, *n* = 10 cells), 5.3 ± 0.5 t^*^ (late migration, *n* = 10 cells), and 5.2 ± 0.6 t^*^ (apical constriction, *n* = 10 cells), with a comparable (≈25%) increase between early and late migration (Fig. 5f).

Simulations on rectangular domains suggest that decreasing actomyosin activity tends to increase the oscillation period independent of domain size (Fig. 5g). To further test the contribution of cell contractility, cell polarity, and cell shape to the cells’ oscillatory behaviour, we changed actomyosin activity and boundary conditions but kept realistic cellular geometries and assessed the oscillation period in the different stages of LEC behaviour (Supplementary Information Sec. S8; Supplementary Fig. S8). We found that applying no flux boundary conditions for each stage and applying different actomyosin activities for the different stages (*ζ* = 40 for early migration, *ζ* = 36 for late migration, *ζ* = 30 for constricting cells), resulted in a straightforward increase in period across the three LEC stages (Supplementary Fig. S8c).

This trend of decreasing period with increasing contractility is not surprising, since we can rewrite the strain-dependent myosin turnover as 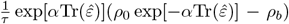. The effective intrinsic timescale associated with this process is 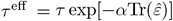, which decreases as the tissue is compressed since 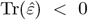 and recalling that *α* = −1. Therefore, as activity *ζ* increases, the intrinsic time scale decreases, and the period of oscillations set by the strain-dependent myosin binding/unbinding also decreases. Thus, it may be surprising that we do not see a further increase in the period as activity is lowered between late migration and apical constriction in Fig. 5. However, we found that keeping the actomyosin activity constant and applying varied boundary conditions for the three stages (weak mixed for early migration, strong mixed for late migration, no flux for constricting cells) caused a drop in the period between late migration (strong mixed boundaries) and apical constriction (no flux boundaries). This means the changing boundary conditions can cancel out the effect of decreasing activity (Supplementary Fig. S8b). When no flux boundary conditions and the same actomyosin activity (*ζ* = 36) were applied to all LEC stages, no significant difference in oscillation period is observed between LEC stages (Supplementary Fig. S8d). Thus, a change in cell size alone does not affect the period.

Overall, our simulations on realistic cellular geometries demonstrate that the observed change of oscillation period between early and late migrating LECs appears to be influenced by actomyosin activity and cell polarity, but not by differences in cell shape.

## III. DISCUSSION

We compared in vivo subcellular patterns of actomyosin contraction of LECs to simulations of a continuum active elastomer model that uses realistic cell geometries and boundary conditions. Notably, we observed distinct contracted regions (CRs), which appear at typically peripheral initiation sites before flowing through the apicomedial network, coalescing into foci and subsequently disassembling. While pulsed contractions in late migrating and apically constricting LECs tend to involve the rhythmical formation and disassembly of a single focus, actomyosin oscillations in early migrating LECs reflect the combined effect of multiple foci in varying stages of assembly and disassembly. We found strong agreement between in vivo data and model simulations in CR number, location, and oscillation period. Interestingly, the model recapitulates the in vivo data best for intermediate levels actomyosin activity that decrease from early to late migration, and again from late migration to constriction. Moreover, cell geometry and boundary conditions need to satisfy two in vivo observations: (1) Cells must round up and shrink over time, and (2) boundary conditions must reflect the change in polarity the LECs undergo, from planar cell polarity during migration to radial cell polarity during apical constriction [14].

The importance of the level of contractility for network dynamics has been shown for LECs [14], *Drosophila* germband cells [11], and amnioserosa cells [54]. That the model predicts a reduction in actomyosin activity in the later stages is unexpected, as junctional cortical tension has been shown to increase when the LECs transit from stationary to migratory behaviour [55], which suggests that pulsating cells increase contractile force during morphogenesis. In addition, our findings highlight the importance of cell geometry in determining the behaviour of actomyosin oscillations. A similar observation has recently been made in in vitro experiments that showed that different contraction patterns can arise solely by changing the system’s geometry [56].

The simple form of active stresses used in the model suggests that the precise form of activity and the complex network of signalling pathways that regulate it is not essential, as long as activity leads to local contractions. Although the continuum model contains no information about molecular mechanisms that regulate actomyosin behaviour in living cells, it can still capture the key behaviours of the actomyosin network. This supports the notion that subcellular actomyosin oscillations are emergent and do not require spatiotemporal heterogeneous upstream signalling [56–59].

Our observation of complex subcellular actomyosin oscillatory patterns in LECs raises the question of their biological significance. For example, these oscillations might help create forces to deform the cell, such as during apical constriction [11, 14, 22]. However, the precise role of pulsed contractions still remains largely unclear [14, 60, 61], as junctional forces alone can drive apical constriction if pulsed contractions are interfered with [4, 14, 62]. On the other hand, in some cases, subcellular actomyosin flows have been shown to be involved in the regulation of cellular processes, such as junctional remodelling [21], cytokinesis [63], embryo polarisation [64], and cell-cell contact formation [65].

Finally, the effects of mechanical cues from neighbouring cells are not understood. It has, however, been shown that they influence the timing of pulsed contractions [22], raising intriguing questions about the synchronisation of oscillations at the tissue scale [61]. Exploring these effect with the current modelling strategy would not be easy, due to the need for deformable domains of integration. Additionally, cellular mechanics are likely driven not just by the actomyosin network but also by cell migration and inter-cellular interactions, making multicellular modelling challenging.

## Supporting information

Supplementary Video Descriptions

## IV. ACKNOWLEDGEMENTS

We thank Chaithanya K.V.S. and M. Rao for many helpful discussions, and A. Iorio and J. Rozman for comments on the manuscript. R.S. thanks D. Mizuno for clarifying aspects of cytoskeleton rheology measurements. We also thank the IT service of the University of Dundee for their support of the HPC infrastructure used in this study. E.D.M. acknowledges funding by the endowed E.N.&M.N. Lindsay PhD studentship. A.B. acknowledges funding from the St Leonard’s College World-leading Doctoral Scholarship, and thanks W.V. Smith for fruitful discussions. R.S. acknowledges support from the UK EPSRC (Award EP/W023946/1).

## V. CONTRIBUTIONS

M.B. and R.S. designed the study. E.D.M. performed analytical and numerical analysis of the model. A.B. performed the experiments and data analysis. J.K. provided supervision in image analysis and interpreted data. J.J. interpreted data. M.B. provided supervision and training in microscopy and image analysis and interpreted data. R.S. provided supervision in modelling and interpreted data. E.D.M., A.B., M.B., and R.S. wrote the paper. All authors reviewed the final manuscript.

## VI. CODE AVAILABILITY

The code used to perform numerical simulations is available at https://github.com/Euan-Mac/Active-Elastomer-Model.

## VII. MATERIALS AND METHODS

### A. In vivo experiments

#### 1. Fly Stocks

Flies were raised on a standard medium. FlyBase [66] entries of the used transgenes are as follows:

*hh.Gal4* : *Scer/Gal4*^*hhGal4*^, *tub.Gal80ts*:

*Scer*\*Gal80*^*ts.αTub84B*^,

*UAS.gmaGFP* :*Moe*^*Scer*\*UAS.T:Avic*\*GFPS65T*^ [44],

*UAS.LifeActin-*

*Ruby* :*Scer*\*ABP140*^*Scer*\*UAS.T:Disc*\*RFPRuby*^, *sqh::GFP* :

*sqh*^*RLC.T:Avic*\*GFP-S65T*^.

#### 2. Expression of transgenes in LECs

*UAS* –transgenes were expressed in the P compartment using *hh.Gal4*. Expression was repressed using a temperature-sensitive allele of the Gal4 repressor, Gal80 (*tub.Gal80ts*), until 30–50 h before imaging. Repression was released by shifting flies from the permissive temperature (25 ^°^C) to the restrictive temperature (29 ^°^C).

Cytoskeletal behaviour is not altered by Gal80ts expression [14].

#### 3. 4D microscopy

Pupae were staged according to [67]. The sex of the experimental pupae was not determined. For imaging, a window was dissected into the pupal case and pupae were mounted in an imaging chamber as described in [68]. All imaged flies developed into pharate adults and the majority hatched. For comparability of individual cells, we recorded LECs at the anterior edge of the P compartment that were neither on the dorsal midline nor in direct contact with histoblasts (Fig. 1a). The same region was analysed in [14]. Z-stacks with two levels and a step size of 0.42–1.14 μm were imaged every 2.7–3.7 s. All images and movies show maximum projections of the two *z*−levels. Imaging was done with a Leica SP8 confocal microscope at 25±2 ^°^C with an HCPL APO CS2 63x/1.40 OIL objective. The image size was 1024×1024 pixels. In all images and movies, the anterior is to the left. Figures and movies were made using ImageJ [69].

### B. Analysis of experimental data

#### 1. Classifying recordings

We focused our analysis on LECs in which contracted regions (CRs) are clearly visible. No analysable samples were excluded; some samples were not analysable, due to drift of the recording or a fold in the pupal cuticle obscuring some of the apicomedial network. LECs in analysable recordings were classified as early migration, late migration, or apical constriction by considering: the extent of histoblast nests at the time of recording; the presence or absence of a lamellipodium; whether the LECs visibly migrate during the recording; and the appearance of the apicomedial actin network, which is more fibrous in late migration compared to early migration.

#### 2. Statistical analyses

Unless otherwise stated in the main text, *n*−numbers for each of the three pulsatile LEC phases, and in total, are as follows: early migration 10 recordings, 8 cells, 8 pupae; late migration – 10 recordings, 9 cells, 8 pupae; apical constriction – 10 recordings, 9 cells, 7 pupae; total – 30 recordings, 26 cells, 23 pupae. The numbers of recordings, cells, and pupae differ because up to two LECs may be recorded in each pupa (one each on the left and right-hand sides of the abdomen), and because some LECs were recorded more than once, where their apical shape and patterns of contracted regions had changed in the intervening time. We performed statistical analyses using Python’s SciPy library, assuming a significance level of 0.05. Box plots were plotted with minimum, first quartile (Q1), median, third quartile (Q3), and maximum. If not stated in figure legends, statistical significance is denoted by the following: \**p <* 0.05, ** *p <* 0.01, and \*\*\**p <* 0.001, where *p* is the *p*−value from the appropriate hypothesis test. Where values are stated in the text, we quote the mean value ± standard deviation of the mean.

#### 3. Quantifying LEC shape

To estimate the lengths of each cell’s long and short axes, we first draw a mask over the cell in every tenth frame of the recording in ImageJ. We then used the Python module Feret [70] to measure the (not necessarily perpendicular) maximum and minimum Feret (calliper) diameters (Supplementary Fig. S2a). The maximum and minimum Feret diameters represent the maximum and minimum distances between two parallel lines tangential to the boundary of the cell mask.

#### 4. Intensity time series

To obtain the intensity time series for each recording, we first draw a mask over the cell in every tenth frame of the recording in ImageJ. Each binary mask was then eroded using a 15 × 15 pixel kernel, to exclude regions close to cell junctions. We then replicated each eroded mask nine times and used the resulting set of masks to extract the mean pixel value within the cell in each recording frame. Since LECs move very little on a timescale of 10 frames (∼ 30 s), applying replicate masks in this manner picked up the majority of signal from the apicomedial actin network, while erosion ensured that fluctuations in the position of cell junctions rarely brought the signal from junctions within the mask. We detrended the time series by subtracting a non-linear least-squares quadratic fit, then denoised using a 10-point cubic Savitsky-Golay filter.

#### 5. Finding peaks and troughs in intensity time series

To estimate the oscillation periods present in the intensity time series, we calculated the time between successive troughs. We first smoothed the detrended time series using a 30-point cubic Savitsky-Golay filter. We then determined the positions of troughs by locating descending zero-crossings of the phase of the corresponding analytic signal, as described in [71]. Here, we used the SciPy implementation of the Hilbert transform. Occasionally, noise in the intensity time series produced two zero-crossings of the phase in close (*<* 70 s) succession. Where this occurred, we discarded the time point with the higher intensity value. We used a similar method of ascending zero-crossings of the phase of the analytic signal to locate peaks.

#### 6. Optical flow and divergence of the velocity fields

We applied CLAHE (Contrast Limited Adaptive Histogram Equalisation) and Gaussian filters to raw images to reduce uneven illumination and noise. We obtained velocity fields using the OpenCV implementation of the Farneback dense optical flow algorithm [72] and filtered velocities using the Gaussian filter to reduce noise, excluding values outside cell masks. We estimated the divergence of the processed velocity fields using a central fourth-order finite difference scheme.

#### 7. Detecting contracted regions

To detect CRs in recordings of LECs expressing GMA-GFP, we first enhanced local contrast using CLAHE filters, then used the aforementioned eroded cell masks (see Sec. VII B 4) to find the mean *μ* and standard deviation *σ* of pixel values within the apicomedial network in each frame of the recording. We then considered a square region of length *l* centred on each masked pixel. If the mean value of all pixels within this region was greater than *μ* + *sσ*, where *s* is a constant, we considered the central pixel to belong to a CR. We selected values of *l* and *s* manually due to varying contrast between recordings, but they all lay in the ranges 3 - 7 μm and 0.1 – 0.9, respectively. After testing all masked pixels, we discarded all detections (i.e. connected components) smaller than 0.5 % of the cell mask area. The remaining detections were manually curated in ImageJ to remove false positives caused by bright vesicles underlying the apicomedial network or individual bright actin fibres. Where no obvious cause of a false positive was noted, we left detections unchanged. An example of detected CRs is shown in Fig. 2e‘ and Supplementary Video 2.

For classifying detected CRs as belonging to either foci or assembling/disassembling foci, we consider a CR to be a focus if it has reached its maximum extent prior to shrinking or fragmenting. At a given time point, multiple CRs which will eventually coalesce are classified as a single assembling focus. Likewise, multiple CRs arising from the fragmentation of a given focus are classified as a single disassembling focus.

As described in the main text, we define initiation sites as locations where distinct CRs were first detected before propagation through the network, focus formation, and disassembly of the focus. Due to contrast limitations, we occasionally did not detect CRs propagating through the apicomedial network in a particular frame. Therefore, to avoid false positives, we manually mapped the locations of initiation sites. We divided cells into nine regions by subdividing the short and long axes into thirds. We then manually recorded the locations of initiation sites by observing raw recordings overlaid with the contours of detected contracted regions. We analysed each recording twice and checked the results for consistency.

### C. Equations of the active elastomer model

At the time scale of a pulsed contraction (∼100 s), it is reasonable to model the actomyosin network as a twodimensional active elastomer [31] for the following reasons. (1) The apicomedial actomyosin network is a quasi-two-dimensional (∼100–200 nm thick and spans the entire cell) highly dynamic network of crosslinked actin filaments and myosin motor proteins that undergo constant turnover [9]. (2) The turnover timescales of actomyosin networks vary but are typically in the range of several seconds to minutes [11, 21, 73, 74]; (3) Micro-rheology experiments have shown that at intermediate time scales – shorter than the time it takes for the cytoskeleton to remodel fully, but longer than other cytoskeletal processes – the actin cytoskeleton exhibits an elastic response [75]. Therefore, we describe CRs in LECs with a continuum active elastomer model [29–32]. We ignore inertia and assume that permeating fluid has no effects beyond providing drag on the actomyosin network moving against it. Under these assumptions, the overdamped dynamics of the network can be described as a force balance between passive and active mechanical forces and friction due to drag [Eq. (1)].

Assuming small strains and an isotropic and homogenous network, we have

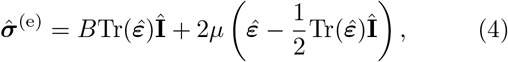

where 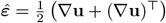, with ⊤ denoting transpose, is the strain tensor, and *B* and *μ* are the bulk and the shear moduli, respectively. Î is the two-dimensional unit tensor. Similarly,

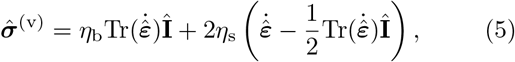

where *η*_b_ and *η*_s_ are, respectively, the bulk and the shear viscosity, and 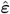 is the strain rate tensor.

#### 1. Dimensionless Equations

Equations (1) and (3) together with constitutive relationships [Eqns. (2), (4), and (5)] were made dimensionless as follows.

We set the unit of length as the screening length 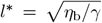, unit of time as a ratio of the square of this length and the myosin diffusion constant, i.e. *t*^*^ = (*l*^*^)^2^*/D*, unit of density as the non-linearity in active stress, i.e. *ρ*^*^ = 1*/ζ*_2_. This allows us to derive the following dimensionless equations,

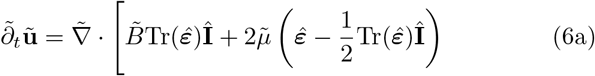

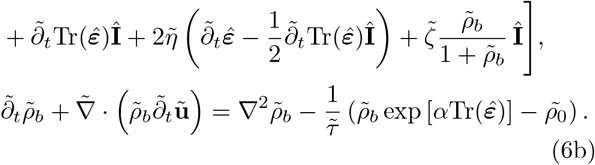

Details of the derivation are given in the Supplementary Information Sec. S3.2. In the text and all figures containing model data, we quoted the dimensionless values of the parameters. There are, however, several free parameters.

To reduce the size of the parameter space to a manageable size, we set *B* = *μ* = 1, *η* = 1, *α* = −1 throughout. We then varied the myosin turnover time scale *τ*, the activity *ζ*, and the preferred myosin density *ρ*_0_. We show all parameters and their values in Supplementary Tab. S1.

We remark that relating the model parameters to the molecular processes is extremely hard to do accurately. Nonetheless, we make rough, order of magnitude estimates. Assuming that *η*_b_ ∼ 2 · 10^−2^ Pa·s [76, 77], and *γ* ∼ *η/ξ*^2^ [45], where *η* is the dynamic viscosity of the cytosol (assumed to be comparable to that of water [78], i.e. *η* ≈ 10^−3^ Pa·s^−1^) and *ξ* ∼ 50 nm is the characteristic actomyosin mesh size [79], we estimate *l*^*^ ∼ 0.25 μm. With the myosin diffusion constant estimated to be *D* ∼ 0.01 μm^2^/s [31], we estimate *t*^*^ ∼ 5 s.

#### 2. Weak Form

We solved Eqns. (6a) and (6b) numerically on domain Ω with boundary *∂*Ω using the finite element method (FEM) implemented in FEniCS [50, 51]. To do this, we recast Eqs. (6a) and (6b) into a weak form as follows

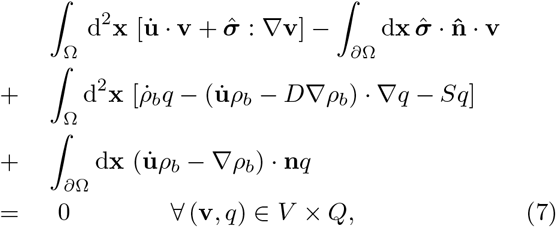

where **v** and *q* are test functions, drawn from the space of test functions *V* and *Q*. The boundary conditions can be applied through the boundary integrals. For example, since throughout this work we used **u** = **0** on *∂*Ω it can be shown that it is also valid to specify that **v** = **0** on *∂*Ω, ∀**v** ∈ *V* [50, 51], meaning that the first boundary integral in Eq. (7) vanishes. For the myosin, we either have a constant myosin boundary condition, which means the boundary integral vanishes by the same argument as for **u**, or we have a no flux myosin boundary condition, meaning 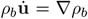, so the second boundary integral in Eq. (7) also vanishes.

We then discretised the system in time using the backward Euler method, giving the following weak form

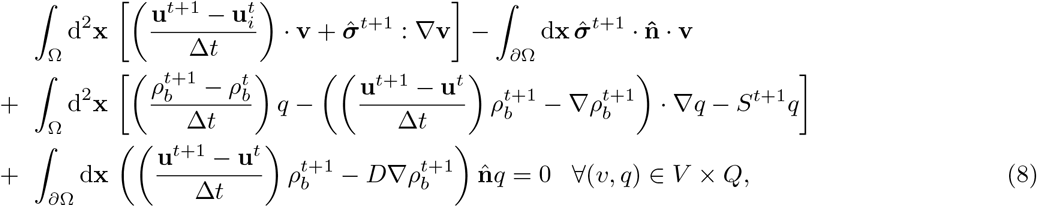

where *t* denotes the time step and Δ*t* denotes the step size in time, 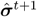, denotes that 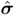 should be evaluated using the values of the fields at *t* + 1 not *t*. These equations result in an implicit system of equations which defines 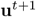 and 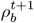 using **u**^*t*^ and 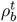.

#### 3. Mesh Construction

We constructed the FEM meshes using the meshing software Gmsh [80]. The general procedure was to first obtain a set of points which form the boundary of the surface under consideration. We then fitted a b-spline curve to these points and from this, we used Gmsh to construct a triangulated surface.

For studies of real cellular geometry, we constructed a cell mask by manually segmenting the first image in a given recording as discussed in Sec. VII B 3. We then used the image analysis library openCV to extract a list of points which form the boundary of this mask [72]. From here we used the standard meshing procedure described above.

### D. Analysis of numerical solutions

#### 1. Tracking of CRs in numerical data

To count the number of CRs observed at each time point, we used the following scheme. First, we computed the mean value and the standard deviation of myosin density across all points in space and time. At each recorded time step, we found all vertices in the FEM mesh where the value of *ρ*_*b*_ was greater than two standard deviations above the mean value. We then ran a standard clustering algorithm based on connected component labelling to identify which of these high myosin vertices were connected by a continuous unbroken path of neighbouring vertices. This gave us a list of high myosin clusters at each time step. Finally, we discarded any clusters made up of a single vertex and labelled the remaining clusters as contracted regions.

To track the motion of a CR, we sought to associate clusters over time. We achieved this by computing the distance between all pairs of vertices between all possible pairs of clusters in consecutive time steps. We then checked if the minimum of the distances was less than a tolerance value, set to 2*l*^*^. If this was the case, we considered the two clusters as the same CR.

The location of the initiation site is important. We split each cell into nine regions distributed as a 3×3 mesh as follows. We first aligned the cell’s long axis to point in the vertical direction. We then identified the vertical location of the initiation site. To find its horizontal location, we then measured the cells’ width at that vertical location and binned the position of the initiation site position accordingly.

#### 2. Finding Peaks and Troughs in Time Series

In several instances, it is useful to be able to locate the peaks in a spatially averaged time series. To do this, we used the SciPy find peaks algorithm. This gave us the location of the local maxima in the time series. Since there were sometimes small localised peaks that did not represent the system’s full oscillations, we also excluded all peak values that were not within the top 20 % of all values in the times series. We used the same procedure to compute the troughs by repeating the process with the sign-flipped time series.

To compute the dominant mode of oscillations in the numerical data, we first computed the periodogram of myosin density at each spatial location on the mesh. This was done using the SciPy library’s periodogram function with the detrend option set to constant to allow the mean signal to be subtracted. We then averaged this periodogram over all points in space and computed the dominant mode as the frequency with the largest amplitude. The period is the reciprocal of this dominant frequency.

## Supplementary Information

## S1 Supplementary figures

**Supplementary Figure S1.**
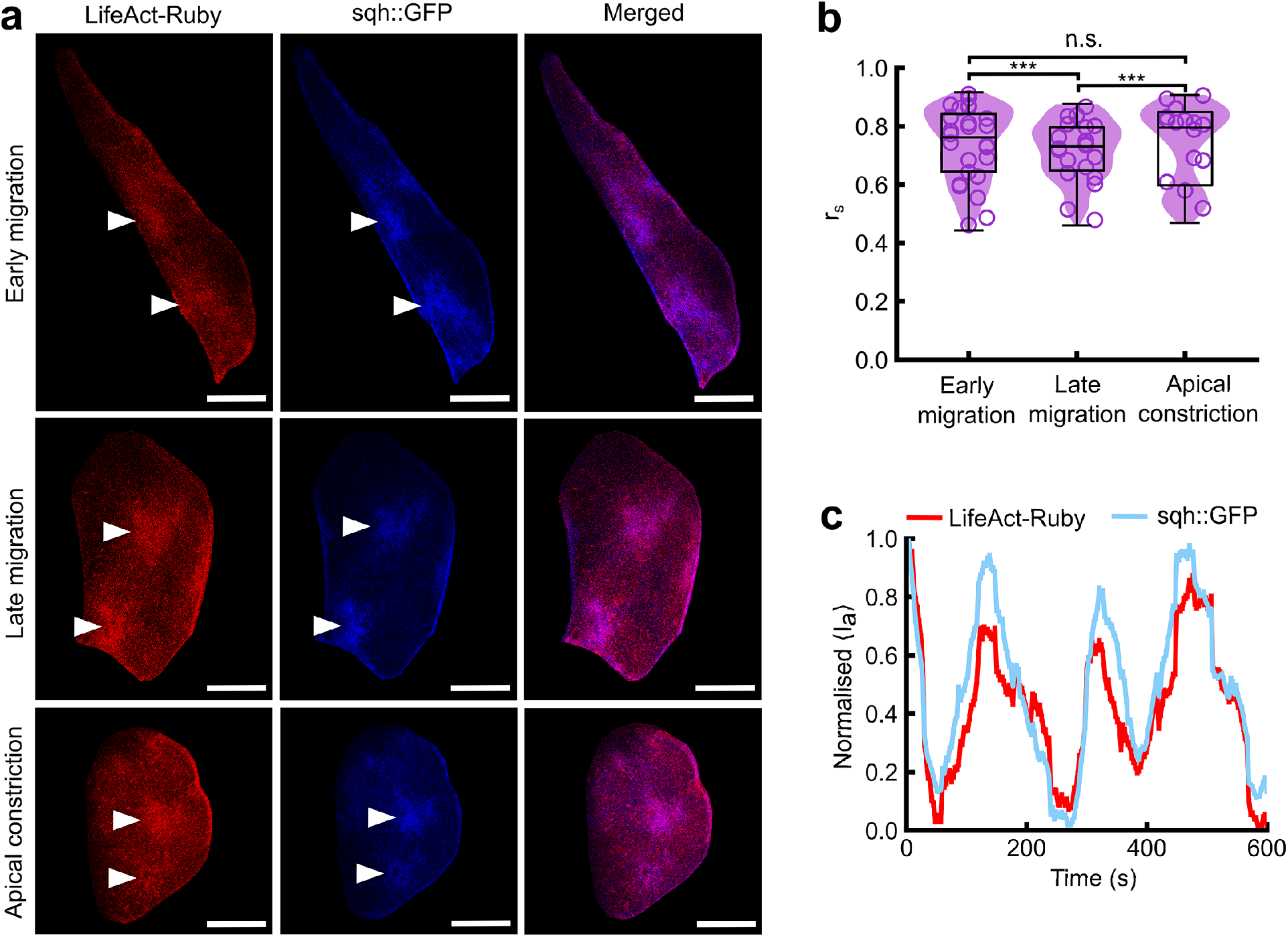
The distribution of F-actin within the apicomedial network is a good proxy for that of myosin II. **a**. LifeAct-Ruby and Sqh::GFP (Myosin Regulatory Light Chain) show similar high-intensity CRs (white arrows) in all three pulsatile LEC stages. All scale bars 20 μm. **b**. Spearman correlation of LifeAct-Ruby and Sqh::GFP fluorescence intensity for each pulsatile LEC stage. A correlation coefficient was calculated at each time point in recordings with a time interval of 3 s and a duration 10 min. Early migration 0.74 ±0.12 (n = 1001 frames, from 5 cells); late migration 0.72 ±0.10 (*n* = 800 frames, 4 cells); apical constriction 0.74 ±0.13 (*n* = 600 frames, 3 cells). *p <* 0.001 for all data points. Images were masked to exclude background and denoised using a Gaussian kernel with a standard deviation of 3 pixels before calculating each correlation coefficient. For clarity, every 40^th^ data point is shown. **c**. Representative time series for spatially averaged apical LifeAct-Ruby and Sqh::GFP fluorescence intensity show similar oscillations. \**p <* 0.05, ** *p <* 0.01, \*\*\**p <* 0.001

**Supplementary Figure S2.**
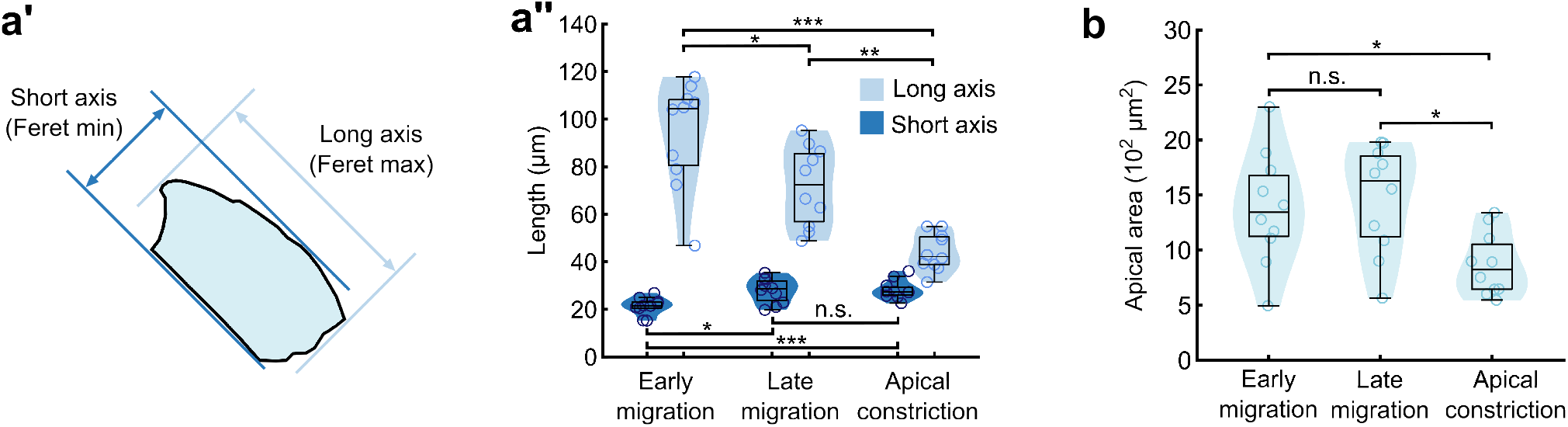
LECs reduce apical aspect ratio and area when transitioning from migration to constriction. **a**’. The long and short axes of each cell’s apical surface were determined by calculating the minimum and maximum Feret diameters, which are the minimum and maximum distances between two parallel lines tangent to the cell boundary. **a**’’. Long and short axis lengths for LECs from early migration to apical constriction show a reduction in apical aspect ratio from early migration to apical constriction. Early migration 21±1 μm, 94±7 μm; late migration 28±2 μm, 72±5 μm; apical constriction, 28±1 μm, 44±2 μm. *n* = 10 recordings per stage. **b**. LEC apical area remains similar from early migration (1380±490 μm^2^) to late migration (1460±470 μm^2^), and decreases significantly as cells enter the constriction phase (870±270 μm^2^). The apical area was measured at 30 s time intervals and averaged over each 10 minute recording. *n* = 10 recordings per stage. \**p <* 0.05, ** *p <* 0.01, \*\*\**p <* 0.001

## S2 Supplementary Videos

### S2.1 Supplementary Video 1

#### LECs undergoing pulsed contractions exhibit dynamic regions of high F-actin density

Confocal micrographs of early migrating, late migrating, and apically constricting LECs expressing the F-actin marker GMA-GFP. Migrating LECs generate a lamellipodium along their posterior edge (yellow arrowheads). Cells in all three pulsatile stages show dynamic regions of high F-actin density (red arrowheads), which we term “contracted regions” (CRs). All scale bars 10 μm. A: anterior; P: posterior; L: lateral; M: medial.

### S2.2 Supplementary Video 2

#### Contracted regions assemble into foci

CRs (outlined in black) detected in an early migration LEC expressing the F-actin marker GMA-GFP are seen to propagate through the apicomedial actin network before forming high fluorescence intensity foci (magenta asterisks) and disassembling. Scale bar 10 μm. A: anterior; P: posterior; L: lateral; M: medial.

### S2.3 Supplementary Video 3

#### At low activities we see a single CR in the model

Numerical simulation of active elastomer model on a rectangular domain of size 8 × 20*l*^*^. We use a no-flux boundary condition. Activity is set to *ζ* = 38 and *k* = 1.1. This corresponds to the region in the phase diagram in Fig. 3 in the main manuscript with the lowest number of CRs. Other parameters are given in Supplementary Table S1. The myosin field is shown in grayscale and the velocity field is shown with red arrows. We see only one contracted region at any given time. The total length of the clip is 200*t*^*^.

### S2.4 Supplementary Video 4

#### At high activities we see many CRs in the model

Numerical simulation of active elastomer model on a rectangular domain of size 8 × 20*l*^*^. We use a no-flux boundary condition. Activity is set to *ζ* = 44 and *k* = 0.1. This corresponds to the “most active” point in the phase diagram shown in Fig. 3 in the main manuscript. Other parameters are given in Supplementary Table S1. The Myosin field is shown in grayscale and the velocity field is shown with red arrows. We see up to six distinct contracted regions at some stages of the video. The total length of clip is 200*t*^*^.

### S2.5 Supplementary Video 5

#### CRs become more regular with periodic boundaries

Numerical simulation of active elastomer model on a rectangular domain of size 8 × 20*l*^*^. We use periodic boundaries. Activity is set to *ζ* = 40 and *k* = 0.9. Other parameters are given in Supplementary Table S1. The Myosin field is shown in grayscale and the velocity field is shown with red arrows. We see an example of the “multiple interacting CRs” state, but with a slightly more well ordered pattern than in the case of no-flux boundaries.

### S2.6 Supplementary Video 6

#### An extended wave state is seen at low activities with periodic boundaries

Numerical simulation of active elastomer model on a rectangular domain of size 8 × 20*l*^*^. We use periodic boundaries. Activity is set to *ζ* = 30 and *k* = 0.3. Other parameters are given in Supplementary Table S1. The Myosin field is shown in grayscale and the velocity field is shown with red arrows. We see an example of the “extended wave” state seen only in periodic boundaries.

### S2.7 Supplementary Video 7

#### Fig. 4 early migration – experiments

Actomyosin oscillations in the early migrating LEC are depicted in Fig. 4 a’’. Scale bar 10 μm. A: anterior; P: posterior; L: lateral; M: medial.

### S2.8 Supplementary Video 8

#### Fig. 4 late migration – experiments

Actomyosin oscillations in the late migrating LEC are depicted in Fig. Fig. 4 c’’. Scale bar 10 μm. A: anterior; P: posterior; L: lateral; M: medial.

### S2.9 Supplementary Video 9

#### Fig. 4 apical constriction – experiments

Actomyosin oscillations in the apically constricting LEC depicted in Fig. 4 c’’. Scale bar 10 μm. A: anterior; P: posterior; L: lateral; M: medial.

### S2.10 Supplementary Video 10

#### Fig. 4 early migration – model

Numerical simulation of active elastomer model of a cell in early migration (same geometry as video 7). We use a weak polar boundary condition on myosin. Activity is set to *ζ* = 40. Other parameters are given in Supplementary Table S1. We observe broadly similar pattern of contracted regions as seen in the real cell. Myosin field showed in grayscale. The total length of the clip is 400*t*^*^.

### S2.11 Supplementary Video 11

#### Fig. 4 late migration – model

Numerical simulation of active elastomer model of a cell in early migration (same geometry as video 8). We use a strong polar boundary condition on myosin. Activity is set to *ζ* = 36. Other parameters are given in Supplementary Table S1. Simulations reveal broadly similar pattern of contracted regions as seen in the real cell. Myosin field showed in grayscale. The total length of the clip is 400*t*^*^.

### S2.12 Supplementary Video 12

#### Fig. 4 apical constriction – model

Numerical simulation of active elastomer model of a cell in apical constriction (same geometry as video 9). We use a no-flux boundary condition on myosin. Activity is set to *ζ* = 30. Other parameters are given in Supplementary Table S1. Simulations show a broadly similar pattern of contracted regions as seen in the real cell. Myosin field is shown in grayscale and the velocity field is shown with red arrows. The total length of the clip in 400*t*^*^.

### S2.13 Supplementary Video 13

#### Comparison of model behaviour with k=0

Numerical simulations of active elastomer model on a rectangular domain of size 8 × 20*l*^*^. We use a no-flux boundary condition. Activity is set to *ζ* = 44 and *k* = 0.0, leading to no myosin turnover in this case. Other parameters are given in Supplementary Table S1. The myosin field is shown in grayscale and the velocity field is shown with red arrows, in this example, velocity has been scaled down to be 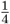 of the real lengths they take on very large values when there is no turnover. We see distinctly different dynamics in the case of no turnover compared with the analogous case but with *k* = 0.25 (supplementary video 4). The total length of the clip is 200*t*^*^.

### S3 Active Elastomer Model

#### S3.1 Model description

**Supplementary Figure S3.**
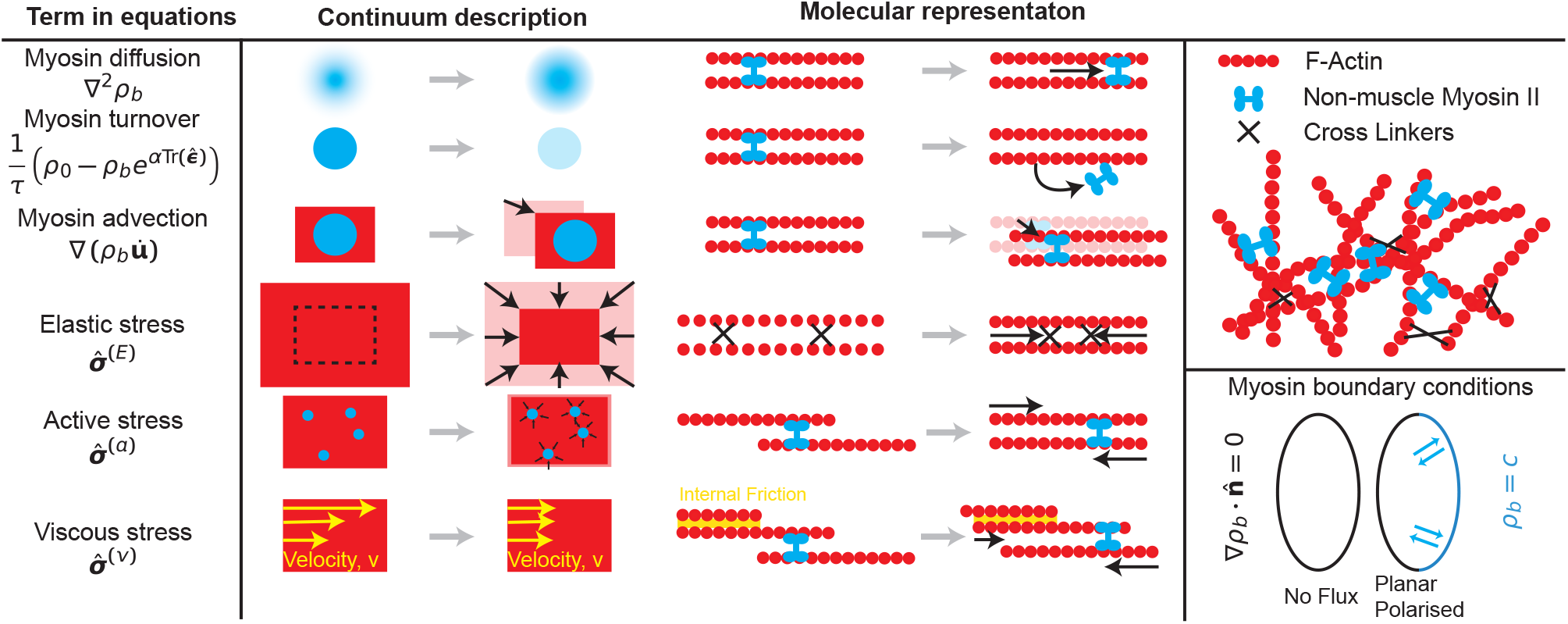
Scheme highlighting the individual parts of the model. The left column shows the key terms included in the equations, along with a pictorial representation of the term, and its molecular origin in the centre column. The top right shows a schematic of the main components included in the model. The bottom right section shows the different possible boundary conditions as described in the main text; light blue represents the part of the cell with the constant myosin condition, representing the lamellipodium.

As discussed in the main text we model the actomyosin network as a two dimensional active elastomer which evolves according to the following (dimensionful) equations

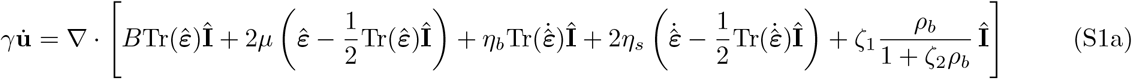

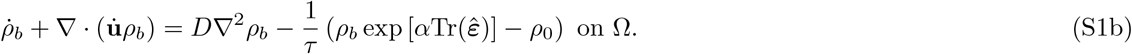

In this section, we give a physical and intuitive picture of the origin of the different terms arising in the active elastomer model. We start with the three contributions to the stress acting on the overall actomyosin network, i.e. elastic, viscous, and active stresses.

Since we are considering processes that occur at time scales short compared to the time it takes the actomyosin network to fully remodel, the network is assumed to behave as a viscoelastic (Kelvin-Voight) solid. To model the elastic response, we introduce the displacement field **u**(**x**), which describes the displacement of a point described by the position vector **x**, i.e. a test particle on the actomyosin cortex placed at **x** is displaced to the position **x**^′^ = **x** + **u**(**x**) due to the deformation of the network. A uniform displacement of the entire network will not lead to the deformation. Therefore, the stress can only arise due to local variations in **u**(**x**). Therefore, the elastic stress is proportional to spatial derivatives of **u**(**x**). If we further assume that the network is homogeneous and isotropic and in the linear regime, this leads to elastic stress which can be written as

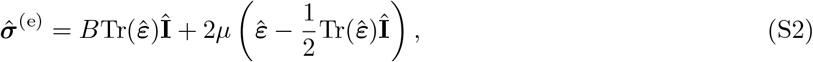

where *B* is the bulk and *μ* is the shear modulus.

The second contribution to the stress is due to viscous dissipation. If we make a reasonable assumption that the local velocity is equal to the time derivative of the displacement vector (i.e. that effects of the nonaffine motion of the network can be neglected), we have 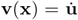. The viscous stress can then be written in terms of the spatial derivatives of 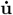, and with the same assumptions of homogeneity and isotropicness. This leads to a viscous stress of the form

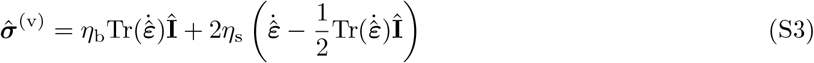

where *η*_*b*_ and *η*_*s*_ are the bulk and shear viscosity of the elastomer.

The last contribution to stress is the active stress due to the action of myosin motors, which we write as

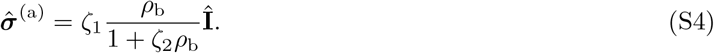

This is a largely phenomenological description of active stress, which comes from the simple assumption that the stress should initially increase with myosin, but must eventually saturate. The activity *ζ*_1_ quantifies the overall amount of stress induced by a given amount of myosin, while *ζ*_2_ quantifies how quickly this effect tails off at large myosin densities.

To derive the equation of motion for the displacement field, we assume that there will be an external drag between the actomyosin network and the cytoplasm, which produces a force per unit volume of 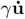, where *γ* is the friction coefficient. We also assume that, since we are interested in a process which occurs over long time scales compared to the molecular processes of the actomyosin network, the system is overdamped, i.e. mechanical forces due to stress remain balanced with the drag force at all times. This leads to Eq. (S1a). The molecular processes driving these three stresses, as well as their continuum equivalent, and mathematical formulation, are represented schematically in Fig. S3.

Since the active stress is dependent on the myosin field, another equation is also required for the evolution of myosin. First, we note that only myosin which is bound to actin filaments generates active stress, so we track only the density of bound myosin density and ignore the unbound myosin. The bound myosin is advected by the motion of the actomyosin network leading to the advection term 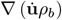. However, myosin motors also move relative to the actin mesh, we describe this process by a diffusion term ∇*D* ^2^*ρ*_*b*_, under the assumption that the actomyosin mesh is isotropic, so there will be no preferred direction of motion.

The myosin can also bind and unbind with some characteristic rates *k*_*u*_ and *k*_*b*_. So a term must be added of the form *S* = *ρ*_*u*_*k*_*b*_ − *ρ*_*b*_*k*_*u*_, however since we assume that the unbound myosin constitutes a bath (i.e. it is present everywhere in approximately equal amounts), this can be written as *S* = *k*_*b*_ − *ρ*_*b*_*k*_*u*_. From previous experimental work [2, 3], it is known that the unbinding rate of myosin can be dependent on the applied strain in an exponential fashion, so we write the unbinding rate as 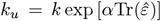]. This leads to a turnover of the following form

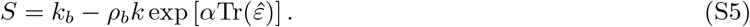

However, this equation is clearer if written in the form

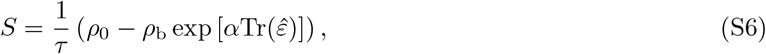

where 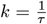 and 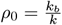, as it becomes evident that we have introduced an intrinsic time scale and a density scale to the equations. These terms, which control the myosin dynamics, are also summarised in table S3.

#### S3.2 Dimensionless Equations

We start with

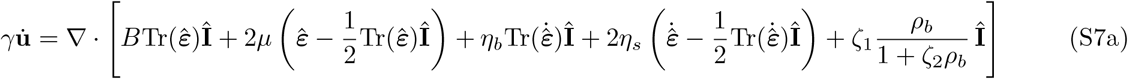

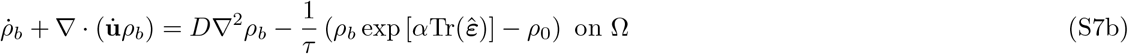

We define the unit of length 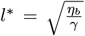, the unit of time 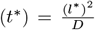, and the unit of density 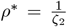. We can then work in terms of the dimensionless variables 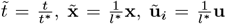, and 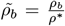. The dimensionless spatial derivatives can then be rewritten as 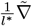, the dimensionless time derivatives are 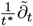.

This means the Eqs. (S7) can be rewritten as

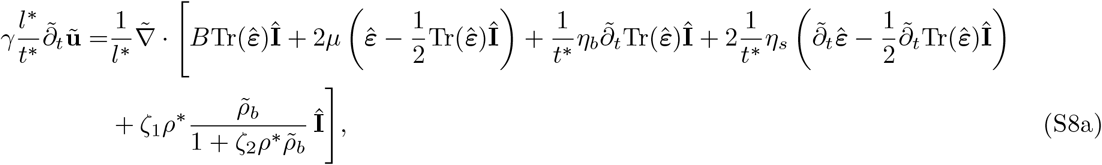

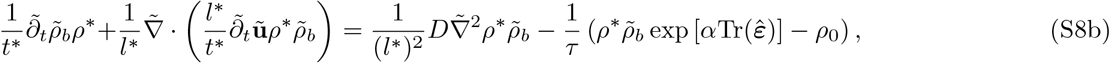

which can be simplified to

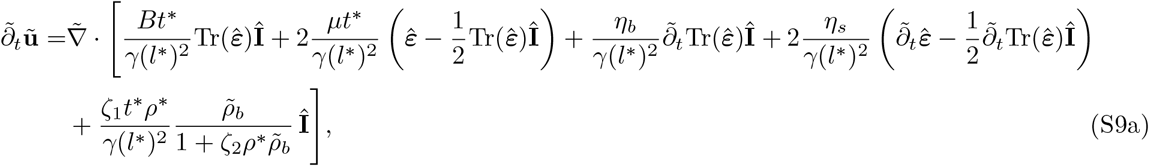

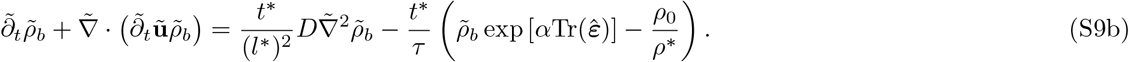

Note that since 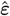 is already dimensionless, we assume 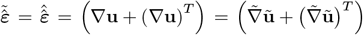. This means the equations are simplified to

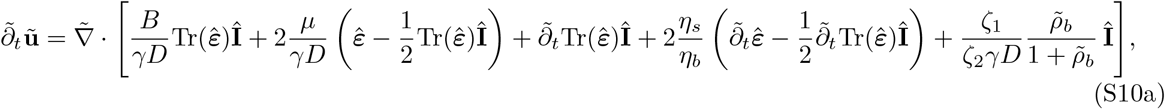

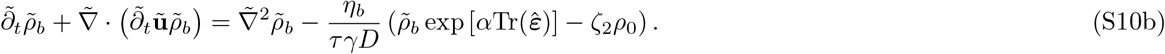

We can then define a new set of constants 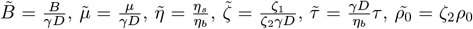. This means the final dimensionless equations are the following

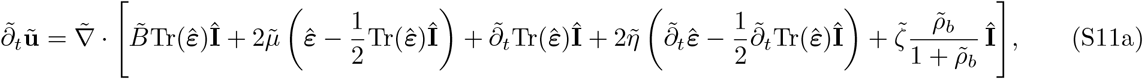

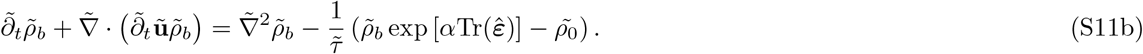

These are the equations that appear throughout the manuscript. For brevity, we drop the tildes, with the understanding that all variables are now dimensionless.

**Table S1.**
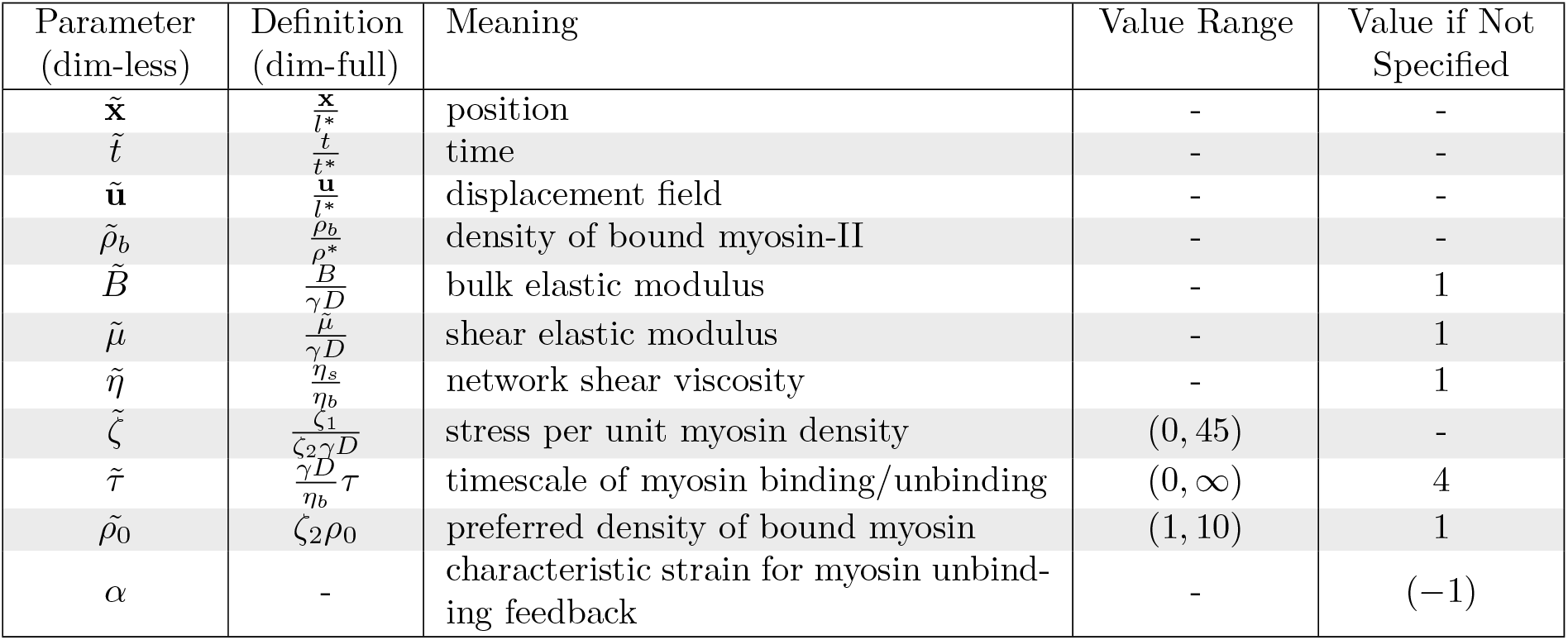
Parameters in the dimensionless equations with their definition in terms of dimension-full parameters and their typical values in used in numerical simulations.

## S4 Simulations Parameters

All simulations were performed with a time step in the range of ∆*t* ∈ (0.01, 0.05) (where the actual value used was adaptive). All meshes were built with a minimum mesh size of 0.5*l*^*^ (using adaptive values). Initial conditions were always set as the homogeneous steady state (*ρ*_*b*_ = *ρ*_0_ and **u** = 0) perturbed by a random Gaussian noise. The noise was rescaled to have values in the range (− 0.005, 0.005). All simulations were run until at least 500*t*^*^, to remove any residual effects of the initial conditions. A typical time series, which shows the evolution to a steady state is shown in Fig. S4a. The dimensionless parameters used in Eqs. (S11) are summarised in Tab. S1 below. To distinguish between dimension-full and dimensionless quantities, we retain tildes on the dimensionless variables.

## S5 Additional Phase Diagrams

In addition, to examining the dependence of the model behaviour on activity *ζ* and the myosin binding/unbinding rate 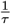, we also examined the dependence on the preferred density of bound myosin, *ρ*_0_, and system aspect ration by tuning length of the long axis *L*_long_. As stated in the main text, our findings suggest that studying the system under periodic boundaries is of limited biological importance, we confined our investigations to the case of natural boundary conditions. The results are briefly discussed in the main text and the phase diagrams are shown here (Fig. S4).

## S6 Mass Conservation vs Myosin Turnover

Note that in the model we include the myosin binding/unbinding effect as a term that appears to violate conservation of mass, since we only consider the bound myosin, and assume the unbound myosin constitutes a bath. This means that if the myosin binding/unbinding time scale gets very large *τ*→ ∞ (i.e. 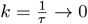), the system becomes mass conserving and the active elastomer becomes a more complex example of the cross-diffusive system recently studied in Ref. [1].

This mass conservation leads to markedly different dynamics, which are clearly visually distinct from the *k* ≠ 0 case, but in a way that is hard to quantify (see Supplementary Movies 13 and 4 for a comparison). One way to understand this is to examine the temporal power spectrum of *ρ*_*b*_. In Fig. S5, we show the temporal power spectrum averaged over space for an elastomer with *k* = 0 (Fig. S5a) and *k* = 0.1 (Fig. S5b) respectively (all other parameters were kept the same). From this, it is clear that by introducing an intrinsic time scale *τ* to the problem, we appear to strongly affect the resonant modes in the system, with all the peaks in the spectrum being at integer multiples of some natural frequency (which is interestingly not exactly equal to *k*) (Fig. S5b). For the case of *k* = 0, however, we see the power spectrum looks much more chaotic, with the peaks no longer being evenly spaced (Fig. S5a). We also compute the highest peak in the power spectrum, for each point on the spatial mesh, and look at the distribution of these dominant frequencies for both cases (Fig. S5c). We can again see that this distribution appears to be much less ordered for *k* = 0 than for the *k* = 0.1 case.

**Supplementary Figure S4.**
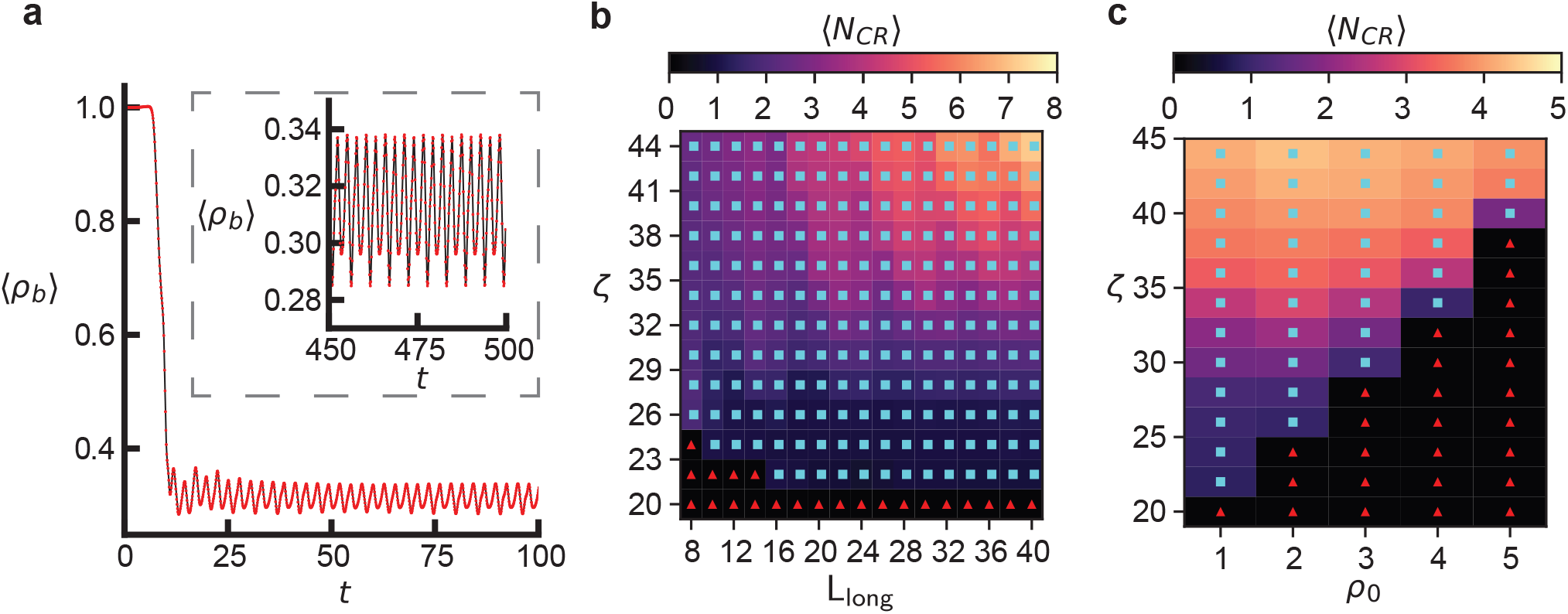
Relaxation to the steady state and additional phase diagrams. **a**. Time series plot of spatially averaged myosin density as a function of time for *τ* = 10 and *ζ* = 44 (other parameters shown in Tab. S1). The system reaches a dynamical steady state after approximately 10*t*^*^. Inset: the same quantity at a much later time showing the steady state is stable. **b**. A phase diagram shows how the average number of contracted regions (CR) depends on the length of the long axis *L*_long_ and activity *ζ*. Size in *y* direction was 8*l*^*^, and 5*L*_long_ × 40 points were used on the mesh. **c**. Phase diagram with *ρ*_0_ and *ζ* being varied. If not specified, parameter values used are given in Tab. S1.

## S7 Linear Stability Analysis

To perform linear stability analysis on the active elastomer model, we start with a slightly generalised version of the full dimensionfull equations [Eqns. (S7)]. To reduce clutter, we use index notation and the Einstein summation convention.

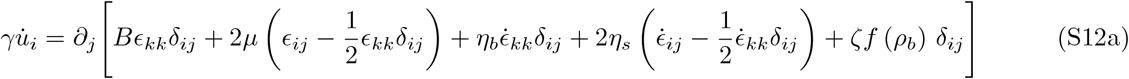

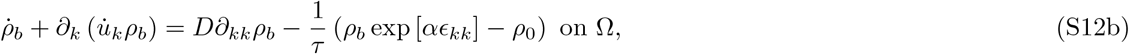

where *i, j* ∈{*x, y*}, *f* (*ρ*_*b*_) is some generalised scalar function of the myosin density and *δ*_*ij*_ = 1 if *i* = *j* and 0 if *i* ≠ *j* is the Kroncker delta symbol. We then linearise the system about its known homogeneous steady state **u** = **u**^*^ and *ρ*_*b*_ = *ρ*_0_. We substitute **u** = **u**^*^ + *δ***u**, *ρ*_*b*_ = *ρ*_0_ + *δρ*_*b*_. By linearising all terms in Eqns. (S12), we arrive at

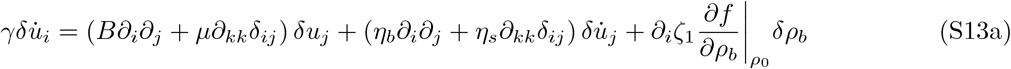

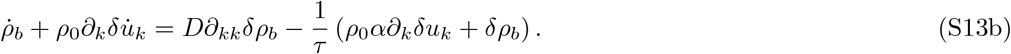

We can then Fourier transform the spatial part, such that 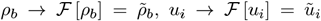, and ∂_*j*_*A* → ℱ [∂_*j*_*A*] = −*iq*_*j*_*Ã*, where *A* is an arbitrary function of position and *Ã* is its Fourier transform. This leads to the following equations in Fourier space

**Supplementary Figure S5.**
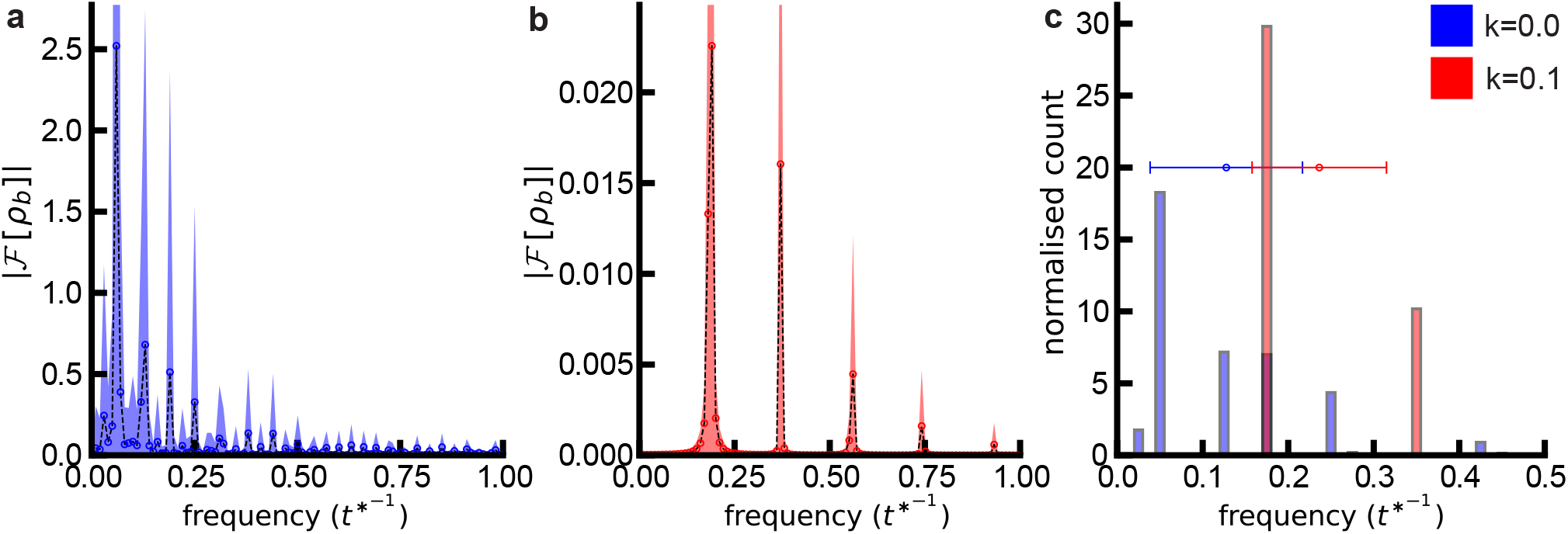
Power Spectrum with and without mass conservation. **a**,**b**. Spatially averaged power spectrum of *ρ*_*b*_ (in frequency domain) with *k* = 0 (**a**) and *k* = 0.1 (**b**). The process is to compute the power spectrum of *ρ*_*b*_ at each point in space, then average over space, and show this average with the shaded area as the standard deviation. **c**. Computation of the dominant frequency at each point in space (as the largest peak in the spectrum). The distribution of these dominant frequencies with the mean dominant frequency is shown as a point with the errorbars showing the spatial standard deviation. Throughout the figure, we see that when *k* = 0.1 (red), the behaviour in frequency space appears much more ordered than when *k* = 0 (blue). This is likely due to the introduction of an intrinsic timescale. Numerical simulations are the same data as from Fig. 3 in the main text, therefore all other parameters are the ones specified in Tab. S1.

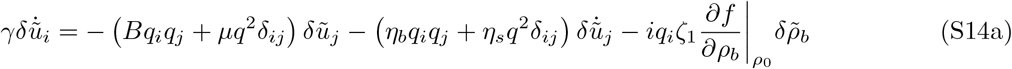

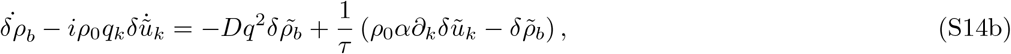

where *q*^2^ = *q*_*k*_*q*_*k*_ = **q** · **q** = |**q**|^2^. These can then be written in matrix-vector form as

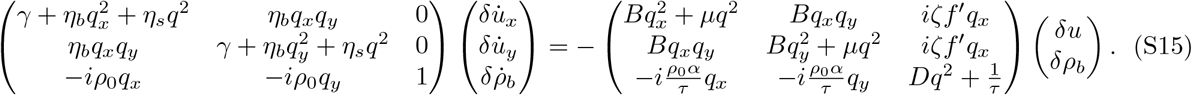

The last expression can be written as

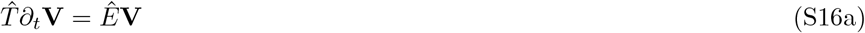

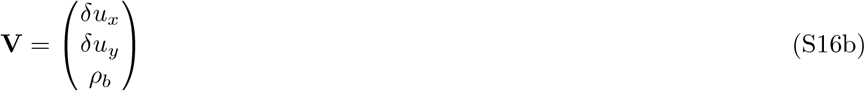

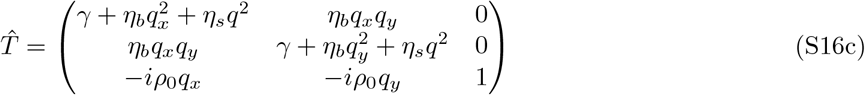

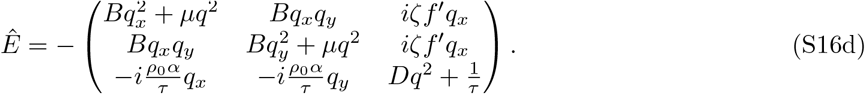

Therefore, we have a generalised eigensystem where the solutions to the set of ordinary differential equations are written as the eigenvectors **w** (*t*) = **w**_0_ exp [*λ* (**q**) *t*], where **w**_0_ is a normal mode formed as a linear combination of *δρ*_*b*_, *δu*_*x*_ and *δu*_*y*_ and *λ*(**q**) are eigenvalues that depend on the wave vector **q**. Finding the stability of the perturbations to the system now becomes a generalised eigenvalue problem of the form 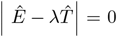. When these equations are solved for *λ*, we find that this gives eigenvalues of the following form

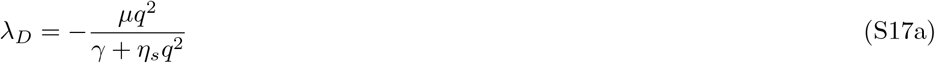

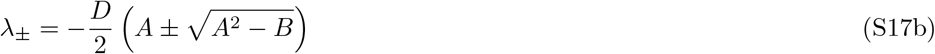

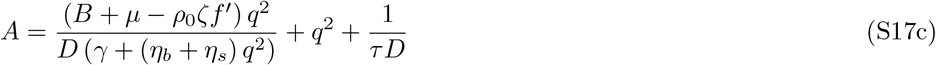

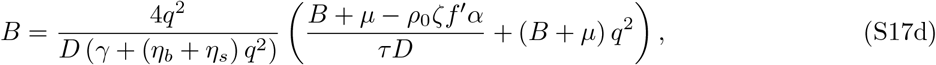

where the first eigenvalue *λ*_*D*_ is a purely diffusive effect, coming from the passive behaviour of the elastomer and the diffusive behaviour of the bound myosin. The other two eigenvalues *λ*_±_, are more interesting and describe the behaviour of the linearised equations.

We can rewrite this in terms of the dimensionless variables as

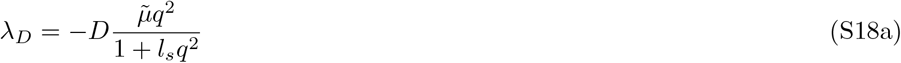

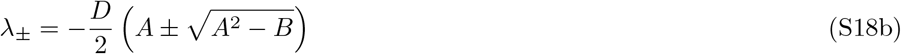

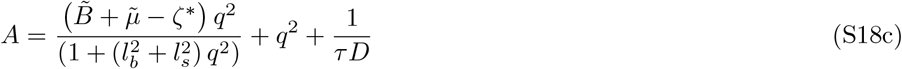

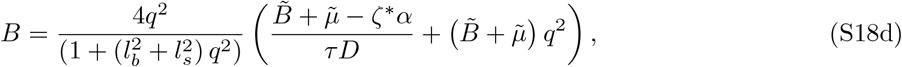

where 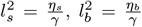 (the previously defined unit of length), *ζ*^*^ = *ρ*_0_*f* ^′^*ζ* which is the analogue of the previously defined dimensionless activity 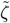, but for the more generalised activity function used here. All other variables are the dimensionless variables defined previously. For brevity, we define the a new slightly modified length scale 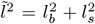 and dimensionless elasticity 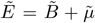 to result in the final dispersion relation

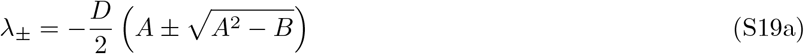

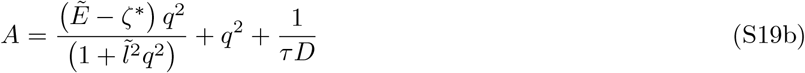

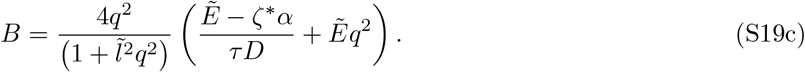

Finally, we can define a dimensionless analogue for the eigenvalue 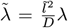, the turnover rate 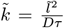, as well as the wavevector 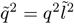 to give a fully dimensionless dispersion relation. Note that these rescalings are intuitively similar but slightly technically different to the ones used in the rest of this manuscript

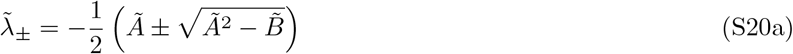

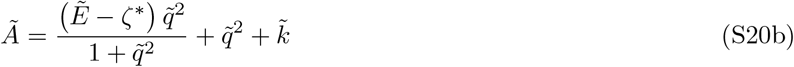

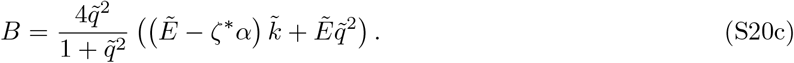

**Supplementary Figure S6.**
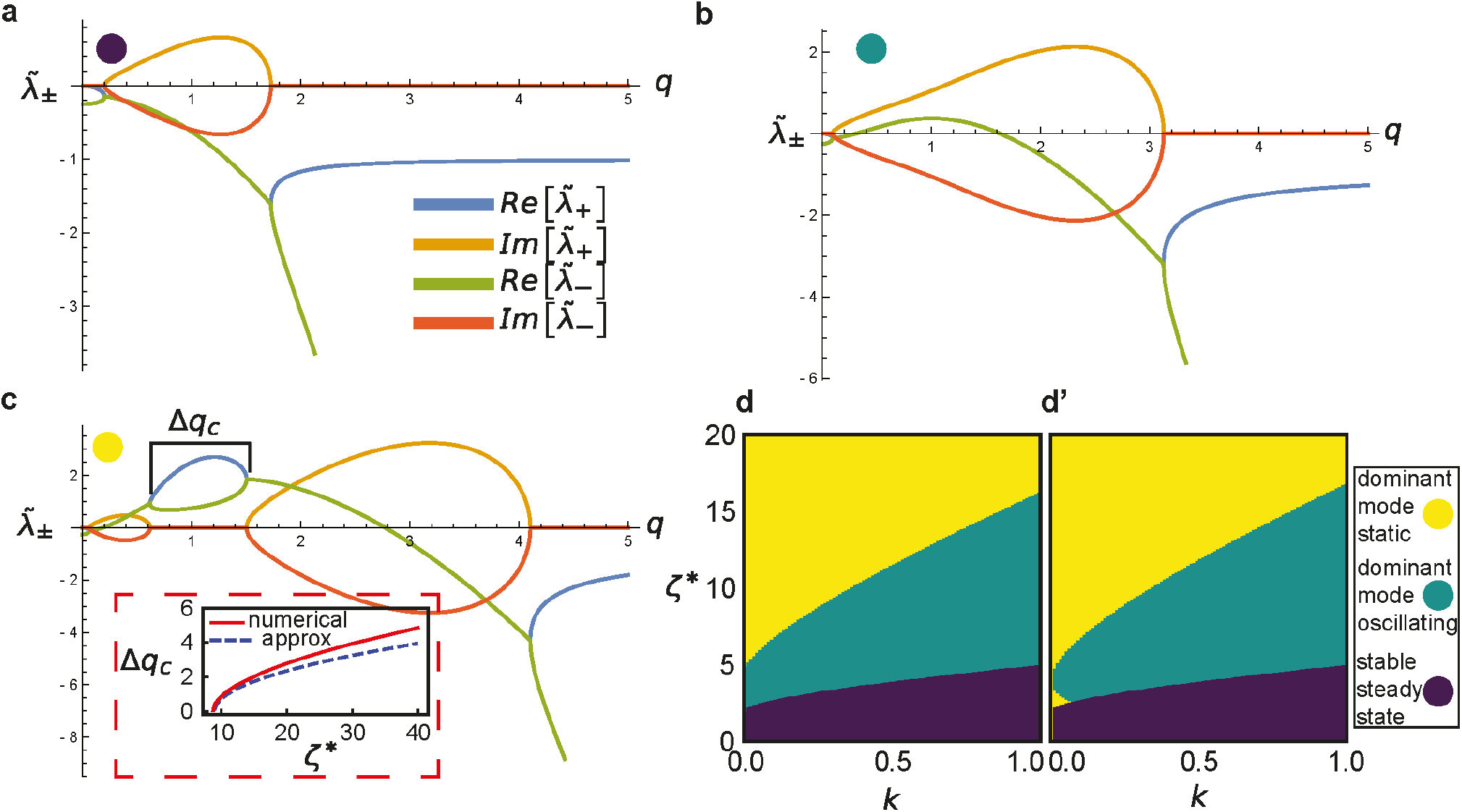
Linear stability analysis. **a-c**. Plots of dispersion relation for linearised active elastomer where 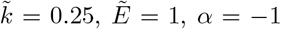. Coloured dots in top left refer to modes shown in d,d’. **a**. When *ζ*^*^ = 1, all modes are decaying, leading to a stable homogeneous steady state. **b**. *ζ*^*^ = 5 leads to some modes (including *q*_max_) with Re [*λ*_+_ (*q*)] *>* 0 and Im [*λ*_+_ (*q*)] ≠ 0, suggesting linearised equations are unstable to oscillatory behaviour. **c**. *ζ*^*^ = 10 leads to modes with both zero and non-zero imaginary parts, however Im [*λ*_+_ (*q*_max_)] = 0, so the expected long-time behaviour is a segregated state. Inset shows width of band of purely non-oscillatory modes as a function of *ζ*^*^ according to the exact dispersion relation, and the approximation derived below. Parameters same as in the other plots. **d**. Phase diagram from numerical analysis of the behaviour of dispersion relation, showing the three identified modes. **d**’. Analytical approximation of phase diagram derived from dispersion relation.

We will drop the tildes on the rescaled variables for the rest of this section. However, we retain the notation *ζ*^*^ to remind the reader that this coefficient is dimensionless but not the same as the dimensionless activity used elsewhere.

To understand the phase behaviour of the system, we consider the behaviour of the fastest growing mode *q*_max_, i.e. the mode which maximises the value of Re [*λ* (*q*)], as this will determine the stability of the homogeneous steady state. If Re [*λ* (*q*_max_)] *>* 0, then the steady state is unstable to perturbations, otherwise the state is stable (since all modes decay). If this is the case, the nature of the instability in the linearised equations is determined by Im [*λ* (*q*_max_)], in particular if Im [*λ* (*q*_max_)] ≠ 0, then the fastest growing mode is oscillatory, so we expect the linearised equations to develop oscillatory dynamics. However, if Im [*λ* (*q*_max_)] = 0 then the fastest growing mode is non-oscillatory, so we expect the linearised equations to develop into a segregated state. To get a better idea of which of these cases apply, in different spaces in parameter space, we plot 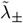 for various parameter values as shown in Fig. S6. From this we see that when the value of *ζ*^*^ is low (Fig. S6a), all modes are decaying in time (Re [*λ* (*q*)] *<* 0), leading to a steady state. As *ζ*^*^ is increased (Fig. S6b), the value of Re [*λ* (*q*)] shifts upwards so that there is some small bands of modes which are growing (Re [*λ* (*q*)] *>* 0) and oscillating (Im [*λ* (*q*)] ≠ 0), leading to the steady state becoming unstable to oscillations. Since the modes are all oscillatory, we know Re [*λ* (*q*)] = *A*. This means that we can compute *q*_max_ as 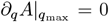, which gives 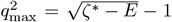. This further implies 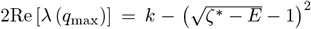. Therefore, the condition for this instability to be reached is that 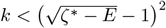.

As the value of *ζ*^*^ is increased further, a small band of the unstable modes switch from having non-zero to zero imaginary parts (Im [*λ* (*q*)] = 0), once this happens it is difficult to analyse the behaviour of the system analytically, as maximising the value of Re [*λ* (*q*)] requires computing the fastest growing oscillatory mode (as we did previously) as well as the fastest growing static mode, and comparing which grows faster. This leads to a algebraically complicated expression, which does not provide much insight. Instead, we note that it always seems to be the case (based on plotting *λ*_±_ for many different values) that once this band of modes with Im [*λ* (*q*)] = 0 appears, the fastest growing mode always appears to lie within them, therefore we expect that once we pass some critical value of *ζ*^*^, and some modes have purely real eigenvalues, we expect one of these modes to dominate at long times, leading to a segregation instability. We can verify this assumption by numerically computing the phase diagram associated to *λ*_±_. We do this by taking a series of vales for *ζ*^*^ and *k* and computing *λ*_±_ for a wide range of evenly spaced values of *q*. Then we simply check which of the set of Re [*λ* (*q*)] values is the largest and use this as the approximate *q*_max_ and therefore categorise the stability of the ready state by checking if Re [*λ* (*q*)] *<* 0 and if it is unstable then we categorise whether the instability is oscillatory or static by checking if Im [*λ* (*q*)] = 0. This leads to the numerical phase diagram shown in Fig. S6d. We find that it is always true that if any unstable mode exists where Im [*λ* (*q*)] = 0 then *q*_max_ always seems to lie within this band as we guessed by the plotting.

Now that we have numerical evidence that the existence of the band of modes with purely real eigenvalues and positive eigenvalues is sufficient to tell us that *q*_*max*_ will lie within this band, we can attempt to construct an analytical expression for when the phase transition occurs that leads to the existence of this band. This band will exist when there exists some *q*_c_ such that *A*(*q*_c_)^2^ − *B*(*q*_c_) *>* 0. Technically, we should also ensure that Re [*λ* (*q*)] *>* 0, which turns out to be equivalent to *A* (*q*_c_) *>* 0) (for *E >* 0, *ζ*^*^>0, *α <* − 1), as long as the previous conditions are also satisfied. This is necessary and sufficient to ensure Re [*λ* (*q*_c_)] *>* 0 and Im [*λ* (*q*_c_)] = 0. Finding an explicit expression for when this inequality is saturated is not particularly easy or insightful, as it involves finding roots of a 4^th^ order polynomial in *q*^2^. Instead, it is helpful to rewrite our main inequality *A*(*q*_*c*_)^2^ − *B*(*q*_*c*_) *>* 0 in the following form

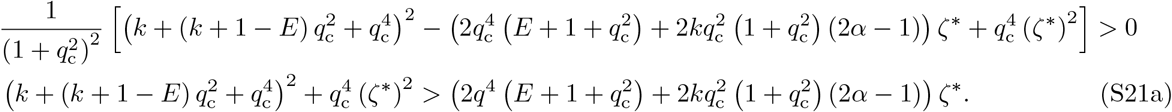

To reduce the complexity of this inequality, we can note two things, first that *k >* 0 and *q*^4^>0 are always true for the cases of interest. Second, since we saw from plotting *λ*_±_ that the band of modes we are interested in typically occurs at values of *ζ*^*^ *>>* 1, the terms of *O*(1) in *ζ*^*^ will be much smaller than the higher order terms in the inequality. The first fact means that if we replace 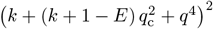 with just (*k* + 1 − *E*)^2^ *q*^4^ in the above inequality, it will remain sufficient but not necessary to guarantee *A*(*q*_c_) − *B*(*q*_c_) *>* 0. The second observation means that since we expect this entire term in the inequality to be small compared to the other two terms, we still expect this bound to be relatively tight. Therefore, we choose to replace Eqn. (S21a) with

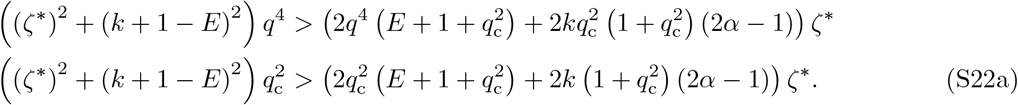

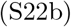

Which finally leads to

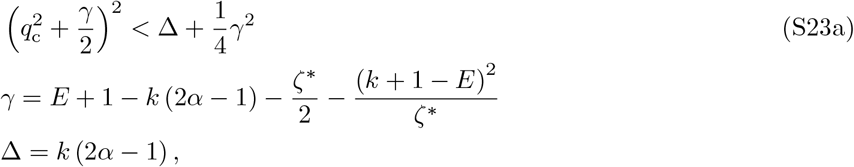

after some algebra. This will have valid solutions when 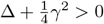, so this final inequality is the required condition for the band of non-oscillatory instabilities to exist, and therefore for the fastest growing mode to be non-oscillatory. We can compute the width of the band of non-oscillatory modes (when it exists) as 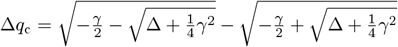. We plot this width according to both the true expression (computed numerically) and the approximate expression derived above in the inset of Fig. S6c, where we see the approximation does fairly well. Although contrary to our expectations, it appears to do better at lower activities.

In summary, we have two main inequalities. The first one is 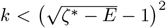, and whenever this is satisfied then the homogeneous steady state is unstable to perturbation. Furthermore, when 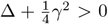 we expect the dominant instability to be non-oscillatory so the linearised equations will be driven towards a segregated state. When this second inequality is not satisfied, we expect it will mostly be the case that the instabilities are oscillatory, however, since the bound is not tight, there are likely non-oscillatory dominant modes close to this boundary. We summarise this analytical estimate of the linear stability phase diagram in Fig. S6d‘ and we can see it matches well with the exact numerical linear phase diagram shown in Fig. S6d.

We note that according to the analysis in this section, our prediction for the behaviour of the active elastomer theory would be that at a low activity we see a stable homogeneous steady state, and as activity is increased this state becomes unstable and the system becomes largely oscillatory, then at even higher activities the system is seen to head towards a segregate state. However, in the numerical solutions from the FEM, we do not see this segregated state at high activity (Fig. 3 in the main text). This surprising observation has been studied in a similar model by Brauns and Marchetti recently [1], and they concluded that this is due to the fact that linear stability analysis about the homogeneous steady state is not a good way to understand the long time behaviour of models such as the one presented here, as they typically lie a long way from this homogeneous steady state. Instead, one should perturb about a non-equilibrium segregated state, and then the prediction of linear stability analysis of the simplified model changes to predict oscillatory behaviour as we saw in the numerics.

## S8 Exploring the Role of Model Parameters

In sections II.E and II.F in the main text, we arrived at the model parameters for each cell stage based on qualitative matching of recordings and simulations. However, it is worth exploring which key parameters control some of the results in Fig. 5 in the main article. To do these we recall that in the simulations shown in Fig. 5 in the main text, three things varied across the simulations of different cells. The first was the cellular geometry, based on the geometry of the real cell being simulated, which is not a free parameter. The other two properties that were adjusted for the cell stage were the values of the actomyosin activity *ζ*, and the boundary conditions. In early migration the cells were assigned *ζ* = 40 with weak mixed boundary conditions, in late migration *ζ* = 36 with strong mixed boundary conditions, and in apical constriction *ζ* = 30 with no-flux boundary conditions. Since these are choices, in this section we investigate three alternative cases. One where *ζ* = 36 (the value corresponding to late migration) for all cell stages, i.e. any observed differences between cell stages must be due to geometry and boundary conditions, one with no-flux boundary conditions for all stages, i.e. any observed differences between cell stages must be due to geometry and activity, and one with *ζ* = 36 and no-flux boundary conditions for all cell stages, so any observed differences must be due to cellular geometry only. The results of these alternative simulations are summarised in Figs. S8 and S7. Note that with these alternative parameters, some numerical simulations fail to show contracted regions entirely, so we exclude these from the analysis.

We found that the correlation observed between cell size, and the mean number of contracted regions, ⟨*N*_*CR*_⟩ (Fig. 5a’’ in the main article) comes from two sources. To understand this, it is instructive to first examine Fig. S7d, where both activity and boundary conditions are kept constant. We see that there is a strong correlation between cell size and ⟨*N*_*CR*_⟩ in the early and late migration cells (dashed line) with a Spearman correlation of *r* = 0.73. However, this correlation becomes weaker when the apical constriction data is included (solid line) with a Spearman correlation of *r* = 0.66. We believe this is because the apical constriction cells were typically much rounder than other cell stages (Fig. S2), which led to the contractile regions typically splitting, instead of having a long axis to flow along, resulting in less orderly dynamics. Since nothing changes in this plot except the mesh used, this correlation is purely an effect of cell geometry.

**Supplementary Figure S7.**
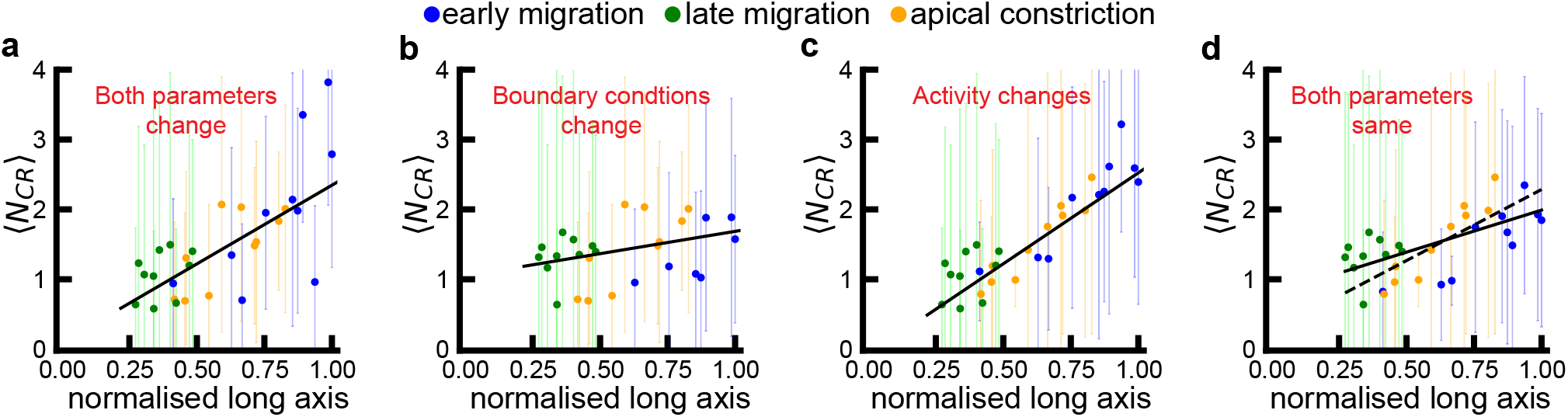
Exploration of the effect of tuning activity *ζ* and boundary conditions on the number of CRs in real cellular geometries. **a**. When both actomyosin activity and boundary conditions change, there is a medium-strength correlation between cell size and the average number of CRs with a Spearman correlation coefficient of *r* = 0.68 with *p* = 0.0002, and a straight line fit with slope *m* = 2.25. The same data as in Fig. 5a’’ is shown for convenience. **b**. When activity is kept constant and only boundary conditions change, there is an insignificant level of correlation between cell size and the average number of CRs with a Spearman correlation coefficient of *r* = 0.36 with *p* = 0.06, and a straight line fit with slope *m* = 0.63. **c**. When boundary conditions are kept constant and activity is varied there is a very strong level of correlation between cell size and average number of contractile regions with a Spearman correlation coefficient of *r* = 0.86 with *p* = 0.0002, and a straight line fit with slope *m* = 2.60. **d**. When activity and boundary conditions are kept the same across all three stages there is a medium strength correlation with a Spearman correlation coefficient of *r* = 0.66 with *p* = 0.0002, and a straight line fit with slope *m* = 1.20 (solid line). However, the correlation becomes even stronger when apical constriction is excluded, leading to a Spearman correlation coefficient of *r* = 0.73 with *p* = 0.0004, and a straight line fit with slope *m* = 2.03 (dashed line). Note in **c**, some cells in early migration failed to show any contractile regions as the activity had been lowered compared to panel **a**, and these were excluded from the analysis. Error bars are all standard deviation. All correlation coefficients were computed with a Spearman correlation test and the *p −* value was computed using a permutation test. The linear fit was done with the Theil-Sen estimator. These were chosen as they are non-parametric methods, and the data was not normally distributed. The long axis is normalised by the length of the longest cell studied.

If we look at Fig. S7b, where the boundary conditions are changed to match the cell stage, we see this correlation becomes much weaker with an insignificant correlation between cell size and ⟨*N*_*CR*_⟩. However, if instead of changing the boundary conditions to match the cell stage, we change the activities and keep all boundary conditions to be no-flux (Fig. S7b), the correlation becomes even stronger with *r* = 0.86. If we change both of these things (Fig. 5a’’ and Fig. S7a), we compromise between these two scenarios and the Spearman coefficient has a value of *r* = 0.68. This correlation between the cell size and number of contractile regions is then partly intrinsic, and partly caused by the changing activities across the three different cell stages. This is likely simply because lower activities cause fewer contractile regions as seen in Fig. 3 in the main text, and the cells in the later stages, which we assign lower activities, are on average smaller than those in earlier stages. The correlation is made less strong by the boundary conditions applied to the different cell stages. In summary we find that the strength of correlations between *N*_*CR*_ and cell length comes from the combined effects of an intrinsic link between these quantities as well as the changing activity *ζ*, linked to cell stages. This correlation is weakened by the changing boundary conditions as the cells change stage.

We also studied the effect of tuning model parameters and boundary conditions on the period of oscillations. We can see that when both activity *ζ* and boundaries conditions are changed, the period of oscillations increases from early migration to late migration and remains approximately constant from late migration to apical constriction (Fig. 5f and Fig. S8a). However, we can see that when both activity and boundary conditions are kept constant, the period does not change at all (Fig. S8d). In addition, we can see that, when only the boundary conditions change and activity is constant, the period decreases when the boundary conditions go from mixed to no-flux (late migration to apical constriction) (Fig. S8b). Moreover, when only activity is changed, this leads to a straightforward increase in period (in agreement with results of main text) (Fig. S8c). Taken together, these results suggest that the changes made to activity cause the period of the system to increase with activity, which leads to the observed period increase between early and late migration (Fig. S8a). However, transitioning from strong mixed to no flux boundaries leads to a decrease in the period – this effect appears to approximately cancel out the increasing effect of actomyosin activity (Fig. S8a).

**Supplementary Figure S8.**
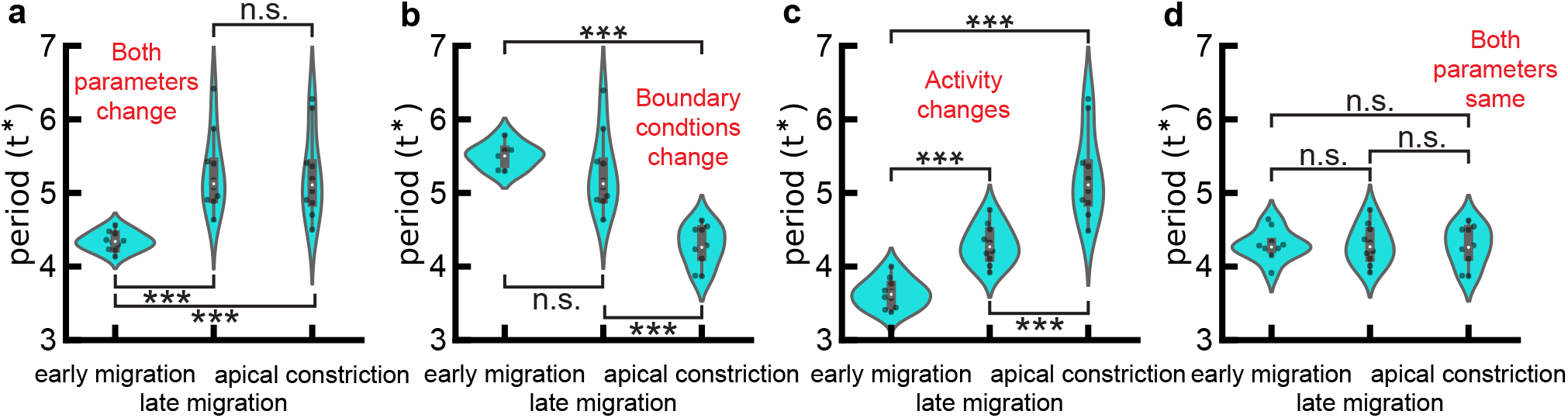
Exploration of effects of actomyosin activity *ζ* and boundary conditions on oscillation periods in real cellular geometries across the three different cell stages. **a**. When both actomyosin activity and boundary conditions change, period increases from early to late migration and then remains approximately constant. Same data as in Fig. 5f in the main text are shown for convenience. **b**. When *ζ* is constant but boundary conditions change, period is seen to decrease as the cells develop from late migration to apical constriction. **c**. When boundary conditions are no-flux throughout the changing activity, a straightforward increase in period is induced. **d**. When only cellular geometry changes we see that this has no effect on the period of the system. \**p <* 0.05, ** *p <* 0.01, \*\*\**p <* 0.001

## S9 CRs Function as Independent Oscillators

In Section II.F.1 in the main text, we hypothesise that contracted regions are independent of each other, leading to interference effects when their effect is combined. To test this hypothesis, we solved the dynamics equations on rectangular domains with natural boundary conditions. We fixed the aspect ratio to 2:1, and set all model parameters to their typical values given in Tab. S1. As the size of the domain increases, the number of CRs also increased (Fig. S9a), consistent with observations in the phase diagram (Supplementary Fig. S4 b). We then measured the “myosin oscillation amplitude” in two ways. (1) We examined each time step and found the maximum and minimum values of myosin across the whole domain (i.e. the cell). We took the ratio of these values and computed the average ratio across all times to get a sense of the average “strength” of the brightest CR in the domain (red plot in Supplementary Fig. S9b). This analysis shows that the larger the domain, the brighter individual CRs become. (2) We averaged the myosin field over space, detected the peaks and troughs of the resultant time series, and measured the peak and trough values in this time series. We then averaged over all peaks or troughs. Then, we took the ratio of this average peak and average trough value (blue plot in Fig. S9b). This analysis shows that the larger the domain, the smaller the amplitude of the oscillations becomes.

Considering all three analyses, we see that both the number of CRs (Fig. S9a) and the “strength” of the brightest CR in the domain (Fig. S9b) increase when the domain size increases, whereas the amplitude of the oscillation decreases (Fig. S9b). This suggests that although CRs are becoming more numerous and individual CRs are getting “brighter”, the combined effect of these CRs is “washed out” due to interference between the CRs, thus, reducing the overall amplitude of the oscillations. This suggests that the CRs are not behaving in a coordinated way, but are interfering with each other, lending evidence to our hypothesis.

We note that the trends observed here are fairly noisy, but this is most likely due to the fact that a lower mesh resolution was used here than in our other analyses, as we needed to be able to use domains which were very large in size, meaning the resolution needed to be reduced to make simulations computationally feasible.

**Supplementary Figure S9.**
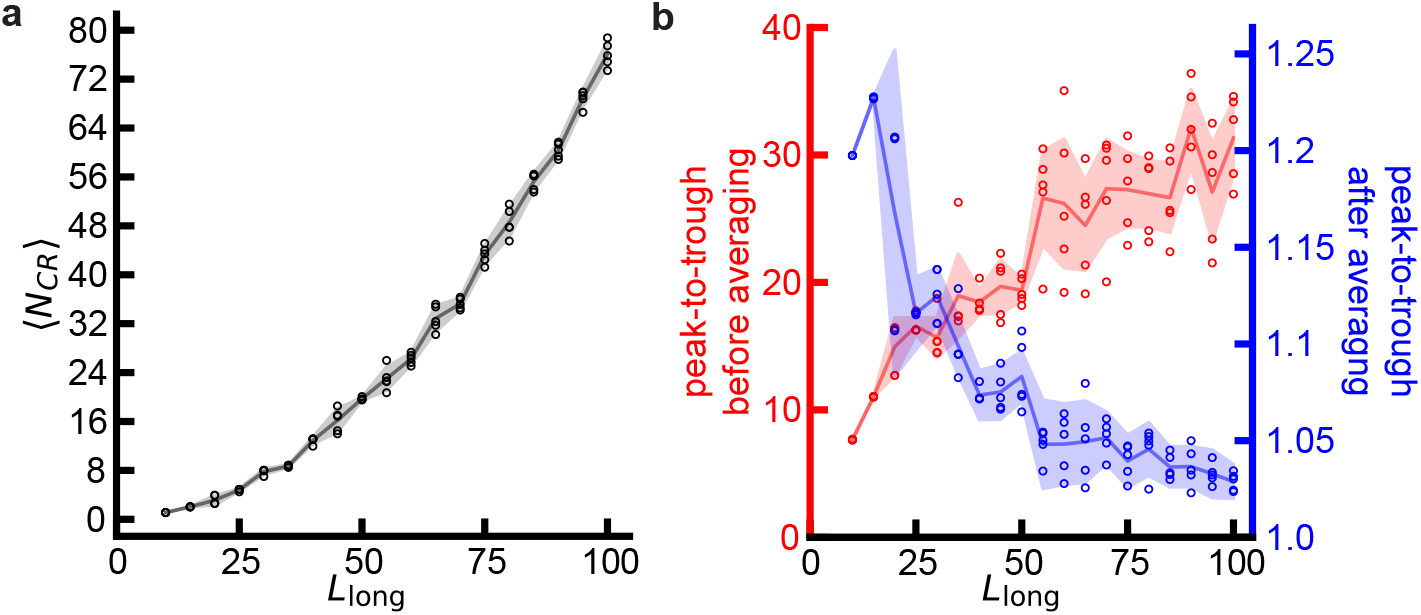
Contracted regions function as independent oscillators. Results from several simulations on a rectangular domain of aspect ratio 2:1, and variable size *L*_*x*_. **a**. The average number of CRs visible at any time is seen to monotonically increase with length of long axis, *L*_long_. **b**. We see that the averaged “strength” of an individual CR, 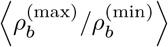, increases with system size, *L*_long_. However, the amplitude of the oscillations in the spatially averaged time series of 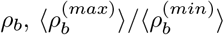 decreases with *L*_long_. Each point represents the mean measurement across one numerical solution. For a given value of *L*_long_, we constructed 5 separate numerical solutions with the same parameters (given in Tab. S1), but different random noise in the initial conditions. Lines represent trends when averaged over all independent simulations with the same *L*_long_, shaded areas show standard deviation. See main text, for full definition of the two types of amplitude measurement plotted.

